# Collagen remodeling promotes a GPCR-mediated mechanosensory immune checkpoint in ADGRG1+ CD8+ T cells and serves as a spatial biomarker of response to immunotherapy

**DOI:** 10.64898/2025.12.19.695209

**Authors:** Sonal Srivastava, John J. Powers, Jay H. Mehta, Sadegh Marzban, Ryunosuke Nozaki, Laxmi S. Karanam, Jacob Torrez, Harish S. Bharambe, Farhoud Faraji, Ritu Chaudhary, Pino Bordignon, Jacqueline Giliberti, Robbert J.C. Slebos, Joseph O. Johnson, Chandler Gantenbee, Xiaofei Song, Marco Cassano, Jose A. Guevara-Patino, Duy T. Nguyen, Alexander R.A. Anderson, Mitsuo Yamauchi, Jeffery West, J. Silvio Gutkind, Christine H. Chung, Antonio L. Amelio

## Abstract

Accessibility of immune cells to the tumor microenvironment (TME) in many solid tumors can be influenced by extracellular matrix (ECM) deposition, organization, and remodeling within tumor stroma. Specifically, lysyl hydroxylase-2 (LH2)-catalyzed lysine hydroxylation in type-I collagen telopeptides leads to formation of intermolecular collagen cross-links creating a stiffened and proteolytically resistant, stable ECM. Advanced head and neck squamous cell carcinomas (HNSCC) frequently exhibit desmoplasia with elevated LH2 expression, and Immune-checkpoint (ICI) therapy is effective in only <20% patients, suggesting that ECM remodeling mechanistically governs the composition and function of TME infiltrates. We show that elevated LH2 and collagen alignment promotes stromal accumulation of CD8⁺ T-cells, and poor response to ICI in HPV-HNSCCs. Integration of clinical biopsies, transcriptomic datasets, and an immunocompetent syngeneic mouse model revealed that aligned collagen spatially restricts adhesion G protein–coupled receptor positive (ADGRG1⁺) CD8⁺ T-cells to activate a non-canonical GPCR-mediated mechanosensory program that drives dysfunction and exhaustion.

**Statement of significance:** Desmoplasia is common in solid tumors, and immune checkpoint inhibitors benefit only some patients. Current biomarkers like PD-L1 and TMB have limited value. Our findings reveal a previously unrecognized collagen-ADGRG1 mechanosensory immune checkpoint, offering a clinically tractable, spatially resolved biomarker to better stratify patients for immunotherapy.

## Introduction

Dysregulated extracellular matrix (ECM) remodeling is a well characterized feature of the tumor microenvironment (TME) that plays pivotal roles in modulating the hallmarks of many solid cancers (1–6). Across anatomical origins and etiologies, fibrotic malignancies display conserved features including hypoxia-driven desmoplasia, aberrant collagen-crosslinking, and stromal reprogramming that shape tumor progression and importantly, immune regulation (7–11). Head and neck squamous cell carcinomas (HNSCC) exemplify highly desmoplastic tumors and provide a clinically relevant context for investigating the immunomodulatory effects of a fibrotic TME (12, 13).

HNSCCs arise from the mucosal epithelium of the oral cavity, oropharynx, hypopharynx, or larynx, and ranks as the sixth most prevalent cancer globally, with about 600,000 new cases annually (14). Established etiological factors include tobacco and alcohol use, as well as human papillomavirus (HPV) infection, with HPV positive HNSCCs exhibiting improved therapeutic responses and survival compared to HPV negative (HPV-) counterparts (15). Current treatment encompasses surgical resection, chemotherapy, radiotherapy, targeted therapy, and immune checkpoint inhibitors (ICIs). Despite the majority of the HNSCCs expressing PD-L1, the overall response rate to anti-PD-1 therapy remains dismal (13-23%) (16–18), and conventional biomarkers such as PD-L1 combined positive score and tumor mutational burden (TMB) provide limited predictive value (19). This knowledge gap emphasizes an urgent need for the development of novel, clinically translatable, tissue-based biomarkers that capture the structural and functional complexity of the fibrotic TME.

Among ECM components, fibrillar collagens govern the biomechanical and biophysical properties of solid tumors (12, 20). Lysyl hydroxylase 2 (LH2), encoded by Procollagen-Lysine, 2-Oxoglutarate 5-Dioxygenase 2 (*PLOD2*), orchestrates collagen stabilization by hydroxylating N- and C-terminal telopeptidyl lysines residues in type I fibrillar collagen prior to cross-linking initiated by lysyl oxidases, promoting the formation of stiff, aligned collagen bundles under hypoxic conditions (21–27). Our previous work demonstrated that *PLOD2*/LH2 is selectively upregulated in late-stage, metastatic HPV-HNSCCs (10) and drives qualitative changes in collagen crosslinking rather than abundance that promotes invasion and metastasis (28). However, the broader implications of this ECM remodeling on immune surveillance and immunotherapy outcomes remain unclear.

Here, leveraging HNSCC as a tractable model, we integrate immune and ECM transcriptomic signatures, clinical biopsies, bulk and single-cell transcriptomic datasets, and a tractable syngeneic orthotopic mouse model, to demonstrate that LH2 is a central regulator of the immune landscape in fibrotic, immunosuppressive TMEs. Using highly multiplexed spatial imaging platforms, we reveal that LH2-mediated collagen stiffening, and alignment reprograms CD8⁺ T-cell organization and promotes dysfunction through mechanosensory immune checkpoint through activation of the adhesion G-protein–coupled receptor ADGRG1 (also known as GPR56).

## Results

### ECM centric molecular profiling identifies distinct TME subtypes in HNSCC

ECM influences tumor development, progression, treatment resistance and immune-cell infiltration (29–31). To investigate the immunomodulatory effects of a fibrotic ECM, we curated an ECM gene signature analysis by curating genes that encompass the major structural stromal elements, receptors that mediate ECM-cell interactions, and enzymes involved in ECM synthesis, remodeling, and degradation, based on published literature **(Table S1)** (32–34). Using this signature, we applied single-sample Gene Set Enrichment Analysis (ssGSEA) to stratify HNSCC samples from TCGA-HNSC (n=515), Moffitt-HNSCC (n=125), and a Moffitt-IO cohort (n=64), into ECM-high and ECM-low groups. Consistent with annotations previously observed for other caners (35), unsupervised hierarchical clustering analysis of immune infiltrates using TIMEx (36) revealed four distinct TME subtypes: Immune enriched-Fibrotic (IE/F), Fibrotic (F), Immune enriched (IE), and Immune depleted (D) **(Figures 1A** and **S1A, S1B)**.

**Figure 1.**
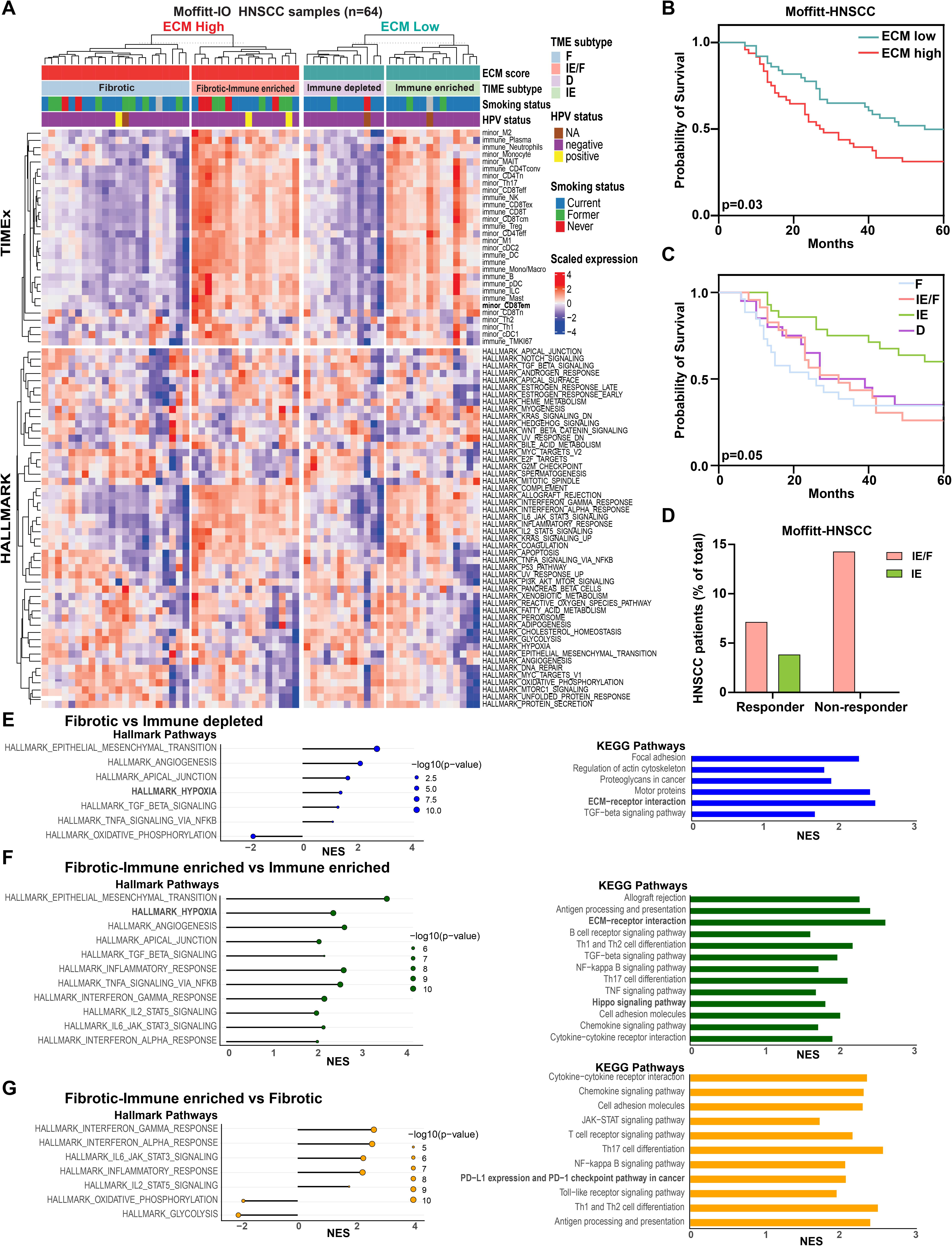
ECM-Immune transcriptomic signature-based TME subtype identification in HNSCC. **(A)** Heatmap classifying TME subtypes in the Moffitt-IO cohort (n = 64) comprising primary treatment naïve HNSCC patients. RNA expression data were z-normalized, each row represents a single gene signature from TIMEx and MSigDB Hallmark Pathways, each column represents a patient sample. **(B)** Kaplan–Meier survival plots for OS of treatment naïve HNSCC cancer patients segregated based on median of ECM score into high (fibrotic) and low (non-fibrotic) for the Moffitt-HNSCC patient cohort (n=125). P values were calculated by Log-rank (Mantel-Cox) test. **(C)** Kaplan–Meier survival plots for OS of treatment naïve HNSCC cancer patients segregated into four different TME subtype for the Moffitt-HNSCC patient cohort (n=125). P values were calculated by Log-rank (Mantel-Cox) test. **(D)** Proportion of HNSCC patients categorized based on response to anti-PD1 therapy into Immune-infiltrated fibrotic (IE/F) and non-fibrotic (IE) subtypes. **(E-G)** Hallmark and KEGG pathways enriched in **(E)** Fibrotic versus Immune depleted subtype **(F)** Fibrotic - Immune enriched vs Immune enriched subtype and **(G)** Fibrotic - Immune enriched vs Fibrotic subtype in Moffitt-IO HNSCC cohort. NES represents the normalized enrichment score, as computed by GSEA; P value <0.05.

The clinical implication of a fibrotic ECM in HPV-HNSCC is demonstrated by a poor 5-year overall survival (OS) probability of the ECM high subgroup across Moffitt-HNSCC and TCGA cohorts **(Figures 1B** and **S1C).** While IE tumors exhibited improved outcomes, the IE/F tumors unexpectedly displayed significantly worse outcomes, comparable to F and D subtypes **(Figures 1C** and **S1D-F).** These results highlight differences between conventional barrier functions that indiscriminately block immune infiltrates in F tumors versus previously unappreciated mechanical features that permit infiltrates but negatively impact immune cell function within IE/F microenvironments, suggesting direct influence on response to immunotherapy. In fact, a retrospective evaluation of nivolumab/pembrolizumab treated patients (Moffitt-IO cohort) showed that despite exhibiting elevated immune infiltration, the IE/F tumors had a higher fraction of non-responders compared to the non-fibrotic IE subtype **(Figure 1D).**

To investigate pathways underlying each TME subtype, we performed Gene Set Enrichment Analysis (GSEA) on a subset of Moffitt-IO cohort with matching tumors on a tissue microarray (TMA), Moffitt-HNSCC, and TCGA-HNSC cohorts using MSigDB Hallmark and KEGG pathways (p-value < 0.05) **(Tables S2-S7)**. Among immune-depleted subtypes (F versus D) hallmark epithelial-to-mesenchymal transition (EMT), hypoxia, and angiogenesis pathways were enriched, along with KEGG TGF-β signaling and ECM-receptor interaction pathways **(Figures 1E** and **S2A, S2B)** consistent with their role of in driving desmoplasia, ECM remodeling, immune exclusion, and tumor progression (37). Comparison of the inflamed subtypes (IE/F versus IE) revealed enrichment of hallmark EMT, hypoxia, TGF-β signaling, and angiogenesis in IE/F subtype, alongside KEGG analysis revealing ECM-receptor interaction, cell adhesion, and TGF-β and Hippo signaling. The IE/F subtype also displayed upregulation of pro-inflammatory hallmark pathways, including IFN-α/γ responses, TNF-α signaling via NFκB, IL-6/STAT3 and IL-2/STAT5 pathways, as well as KEGG chemokine signaling, cytokine-cytokine receptor interaction, B-cell receptor signaling, and antigen processing and presentation indicating the presence of an active immune milieu **(Figures 1F** and **S2C, S2D).**

Within fibrotic tumors, comparison of IE/F versus F subtypes revealed enrichment of hallmark IFN-α/γ responses, IL-6/STAT3, and IL-2/STAT5 pathways but negative enrichment of glycolysis and oxidative phosphorylation. KEGG analysis highlighted antigen processing and presentation, PD-L1/PD-1 checkpoint pathway in cancer, cytokine-cytokine receptor interaction, cell adhesion molecules, chemokine signaling, T-cell receptor signaling, and Th1/Th2/Th17 cell differentiation **(Figures 1G** and **S2E, S2F)**. Together, these findings support that IE/F tumors permit immune cell infiltration, but these cells are functionally impaired by a highly immunosuppressive, fibrotic microenvironment, raising the possibility that ECM architecture directly modulates immune activation, exhaustion, and suppression.

### LH2 expression promotes stiffened collagen alignment and stabilization

To explore the regulators of these transcriptionally distinct phenotypes, differentially expressed genes were compared between ECM-low and ECM-high samples. The *PLOD2* gene encoding the key rate-limiting enzyme LH2, emerged as being significantly upregulated in ECM-high tumors **(Figures 2A** and **S3A-C)**. To validate LH2 expression, 112 tumor cores were stained and scored (0–3) for LH2-positive cell density. Thirty-three cores (29.5%) showed low expression (scores 0–1), while 79 cores (70.5%) showed high LH2 expression (scores 2–3) **(Figures 2B, 2C** and **Table S8**). These tissue cores were then analyzed for collagen features. Picrosirius Red (PSR) staining showed no difference in collagen deposition between LH2-high and -low groups **(Figure 2D, 2E).** Under polarized light, LH2-high cores exhibit increased orange-bright red birefringence, indicating thicker, more mature collagen fibers, whereas LH2-low cores showed more greenish-yellow birefringence, suggestive of thinner, newly deposited collagen **(Figure 2F).** Second harmonic generation (SHG) multiphoton imaging provided label-free, high-resolution maps of collagen architecture used for fiber analyses performed with CT-FIRE, CurveAlign, and OrientationJ **(Figure 2G).** LH2-high tumors contain longer, thicker, and more highly aligned fibers with lower angular variance and increased kurtosis, whereas LH2-low samples displayed shorter, less organized fibers **(Figure 2H-2O)**.

**Figure 2.**
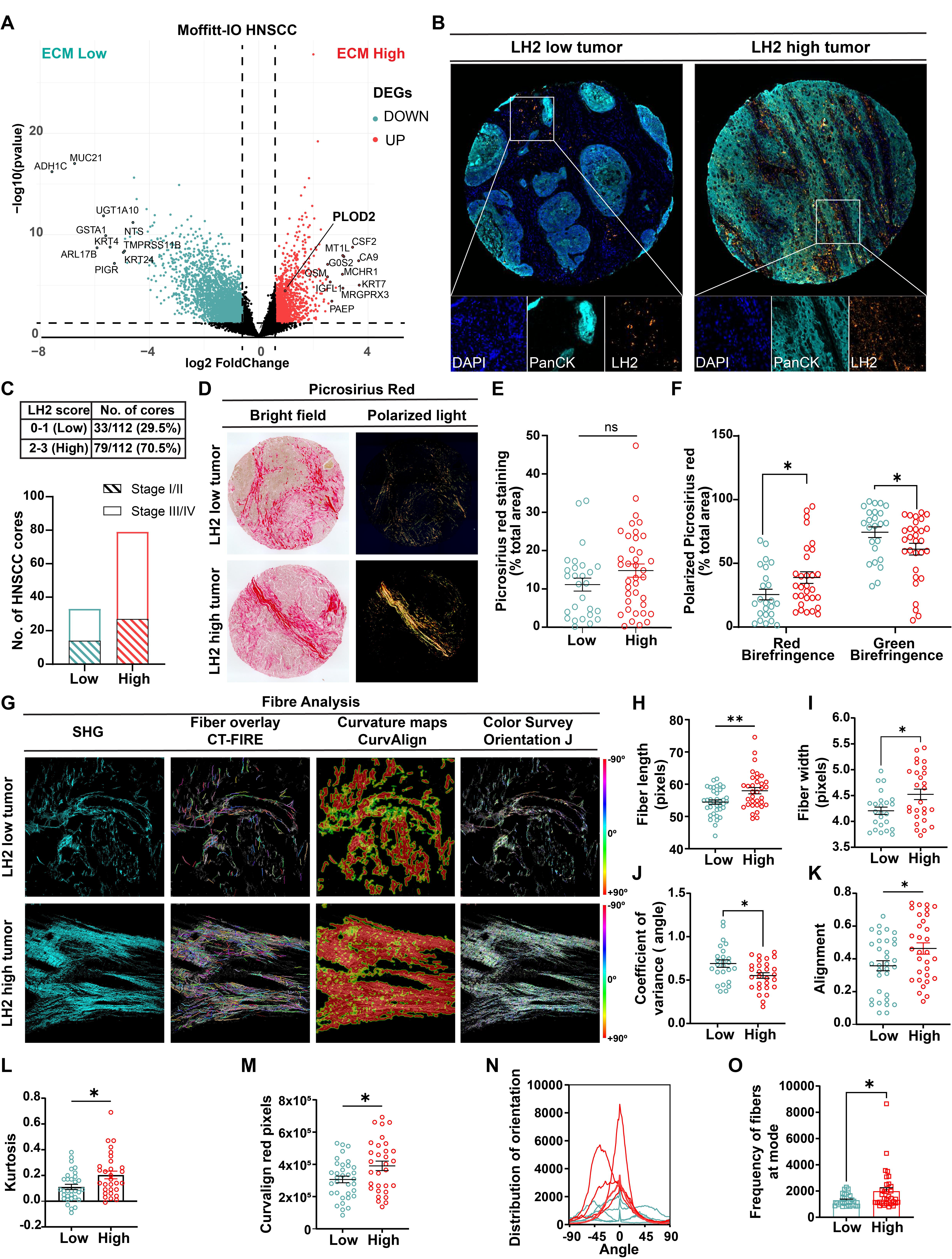
LH2 Expression Drives Collagen Alignment in HNSCC. **(A)** Volcano plot showing differentially expressed genes in ECM-high tumor samples from the Moffitt-IO HNSCC tissue microarray (TMA). **(B)** Representative images of low (n=33) and high (n=79) density LH2 staining in HNSCC cores. **(C)** Distribution of LH2 density scores in HNSCC cores; also represented across early and late stages. **(D)** Representative images of Picrosirius Red (PSR) stained cores visualized under bright field and polarized light. **(E)** Quantification of total collagen deposition in LH2-high (n=28) and LH2-low (n=24) primary tumor cores. **(F)** Quantification of collagen birefringence signals in LH2-high and LH2-low primary tumor cores. **(G)** Representative SHG ROI images of LH2 low and high tumor cores for analyzing collagen fiber features. **(H-J)** Analysis of fiber length, width and angular variance in LH2-low and high tumor cores by CT-FIRE. **(K-M)** Analysis of alignment, kurtosis and red pixel intensity in LH2-low and high tumor cores by CurveAlign. **(N)** Representative Orientation-J plots showing the distribution of collagen fiber orientations in LH2-low (cyan) and LH2-high (red) tumor cores (n = 5 per group). **(O)** Quantification of dominant fiber orientation (mode) frequency across all samples, illustrating the degree of collagen alignment in LH2-low and LH2-high groups. Statistical analysis was performed using two-tailed Mann–Whitney U test. Each dot in the scatter plot represents individual ROI/core, error bars are mean ± SEM. ns, not significant; *P < 0.05, **P < 0.01.

### LH2-modified collagen fibers direct spatial organization of CD3+ T-cell infiltrates

T-cells have been observed to move in parallel to collagen fibers (38), and our prior computational modeling revealed a strong anti-correlation between immune infiltration and disease stage (39). We therefore examined how LH2-mediated type I collagen modifications influence the TME in HNSCC. Multiplex immunofluorescence (mIF) identified CD3⁺ T-cells, CD14⁺ monocyte/macrophage-lineage cells, CD11c⁺ dendritic cells (DC), and Pan-cytokeratin (PCK+) tumor cells (**Figures 3A** and **S4A**). Cell segmentation revealed that LH2-high tumor cores expressed an increased proportion of CD3+ cells (11.1%) as compared to LH2 low cores (7.1%) **(Figure 3B**). However, when these were compartmentalized into tumor (PCK+) and stroma (PCK−) regions, a significantly higher number of T-cells were found to be spatially restricted to the stroma in LH2-high cores (**Figure 3C**). CD14+ cell proportions were higher in LH2-high samples but were similarly distributed across compartments **(Figure S4B)**, while CD11c+ cells did not differ between groups **(Figure S4C).**

**Figure 3.**
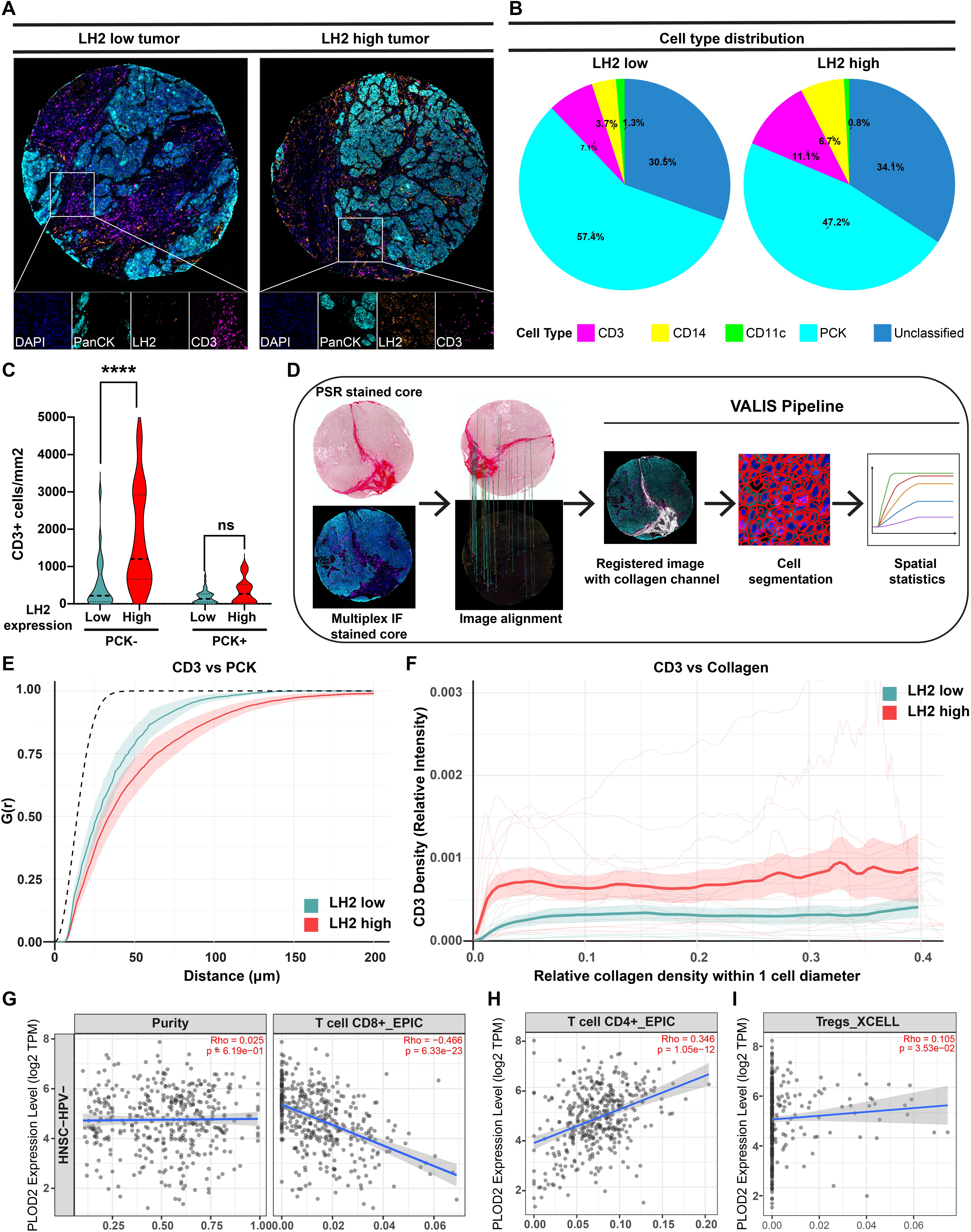
LH2 mediated collagen alignment restricts CD3+ T-cells infiltration in the TME. **(A)** Representative images of tumor cores stained for LH2 (orange), CD3 (magenta), Pan-cytokeratin (cyan) and DAPI (blue). **(B)** Cell types defined by multiplex immunofluorescence and segmentation; pie charts show distributions in LH2-low vs. LH2-high tumor cores. **(C)** Distribution of CD3+ T-cells across tumor (PCK+) and stromal areas (PCK-) of LH2 low (n=25) and LH2 high (n=31) cores. Statistical analysis was performed using two-tailed Mann–Whitney U-test ****P < 0.0001. **(D)** Schematic illustration of the Virtual Alignment of pathoLogy Image Series (VALIS) pipeline. **(E)** Gcross analysis of CD3+ to PCK+ proximity in LH2-high (n=13) and LH2-low (n=11) HNSCC patients. Lines represent group means with shaded areas indicating ±1 standard error. The dashed line denotes spatial independence. Higher curves (LH2 low group) suggest greater association between T-cells and tumor cells. **(F)** Resource selection analysis using the p’’ (rou-hat) rho-hat function showing the relative intensity of CD3+ cells as a function of local collagen density (within one cell diameter), which differs between LH2-high and LH2-low tumors. **(G-H)** Correlation of CD8+ and CD4+ T-cell infiltration with *PLOD2* gene expression in HNSCC TCGA samples using EPIC deconvolution algorithm, tumor purity is depicted on the left. **(I)** Correlation of Treg cell infiltration with *PLOD2* gene expression in HNSCC TCGA samples using X-CELL deconvolution algorithm. TIMER plots indicate computed Spearman’s rho and p-values.

To further examine spatial organization, we built an image-registration pipeline using VALIS (40) to map the mIF-defined cell coordinates onto PSR-stained collagen architecture (**Figure 3D**). Gcross analysis quantified CD3+ T-cell proximity to PCK+ tumor cells, which estimates the probability that a randomly selected CD3+ cell has at least one PCK+ neighbor within a distance r **(Figures 3E)**. At small r values, few CD3+ cells are near PCK+ tumor cells, whereas increasing r reflects greater likelihood of spatial interaction between CD3+ and PCK+ cells. The dotted line in the plot represents the theoretical expectation under a homogenous Poisson distribution, which models complete spatial randomness. In LH2-high tumors, CD3⁺ cells were generally farther from PCK⁺ tumor cells, while in LH2-low tumors, they exhibited closer proximity **(Figures 3E** and **S4D, S4E)**.

Next, the p’’ (rou-hat) resource selection function assesses collagen density as a spatial covariate for CD3+ and PCK+ cell density. Our analysis shows the collagen density differentially influenced CD3+ cell localization **(Figure 3F),** where CD3⁺ cells are more prevalent in collagen-dense regions. In contrast, CD3⁺ cells showed minimal collagen association in LH2-low tumors. PCK⁺ tumor cells exhibited no such differences between these two groups **(Figure S4F)**. These findings suggest that although LH2-high tumors are infiltrated by T-cells, the presence of highly aligned collagen bundles may re-direct these cells within the TME to the stroma and limit their access to, and cytotoxic effects on, tumor cells.

Since CD3 is a pan-T cell marker, we next examined T-cell subtypes using TIMER2.0 to analyze deconvolved TCGA data from HPV-HNSCC data. Immune cell proportions estimated using two independent algorithms, EPIC and TIMER, consistently revealed a significant negative correlation between CD8+ T-cell infiltration and *PLOD2* expression **(Figures 3G** and **S4G).** In contrast, CD4+ T-cells positively correlated with *PLOD2* expression **(Figure 3H** and **S4H),** whereas Tregs **(Figure 3I)** and NKT-cells **(Figure S4I)** showed no correlation with *PLOD2* expression. Amongst the myeloid populations, CD14+ monocyte/macrophage infiltration **(Figure S4J)** and myeloid-derived suppressor cells (MDSCs) positively correlated with PLOD2 expression levels **(Figure S4K),** while CD11c+ DCs showed weak positive correlation **(Figure S4L).**

### Collagen fibers aligned and stabilized by LH2 promote CD8+ T-cell dysfunction

LH2 expression has been associated with advanced-stage HNSCCs and shown to promote metastasis in an immunocompromised orthotopic tongue xenograft model (10, 28). To assess LH2 function in an immunocompetent setting, we established tongue tumors in C57/BL6 mice using 4MOSC1 mouse oral squamous cell carcinoma cells, which exhibit a tobacco carcinogen-related mutanome reflective of HPV-negative, smoking-related HNSCCs (41). We engineered 4MOSC1 cells to express LH2 (TRE3G-*PLOD2*) under a third-generation doxycycline (Dox) inducible promoter along with a constitutively expressing LumiFluor, GpNLuc (42, 43). Dox induced PLOD2/LH2 expression was confirmed at both RNA and protein levels in these stably transduced cells **(Figure S5A-C).**

Tumor burden assessed by BLI following orthotopic transplantation of 4MOSC1-TRE3G-*PLOD2* cells into tongue of C57Bl/6 mice fed either normal or Dox chow (**Figure S5D**) and showed no difference in growth kinetics between Dox and control (No Dox) groups **(Figure S5E, S5F)**, consistent with previous studies using immune-deficient hosts (28). Tumors exhibited typical HNSCC histology **(Figure 4A)** and metastatic progression to draining lymph nodes in the LH2-high group **(Figure S5G, S5H)** as we previously reported but also supporting a role for LH2-modified fibrillar collagen bundles in directing immune suppression.

**Figure 4.**
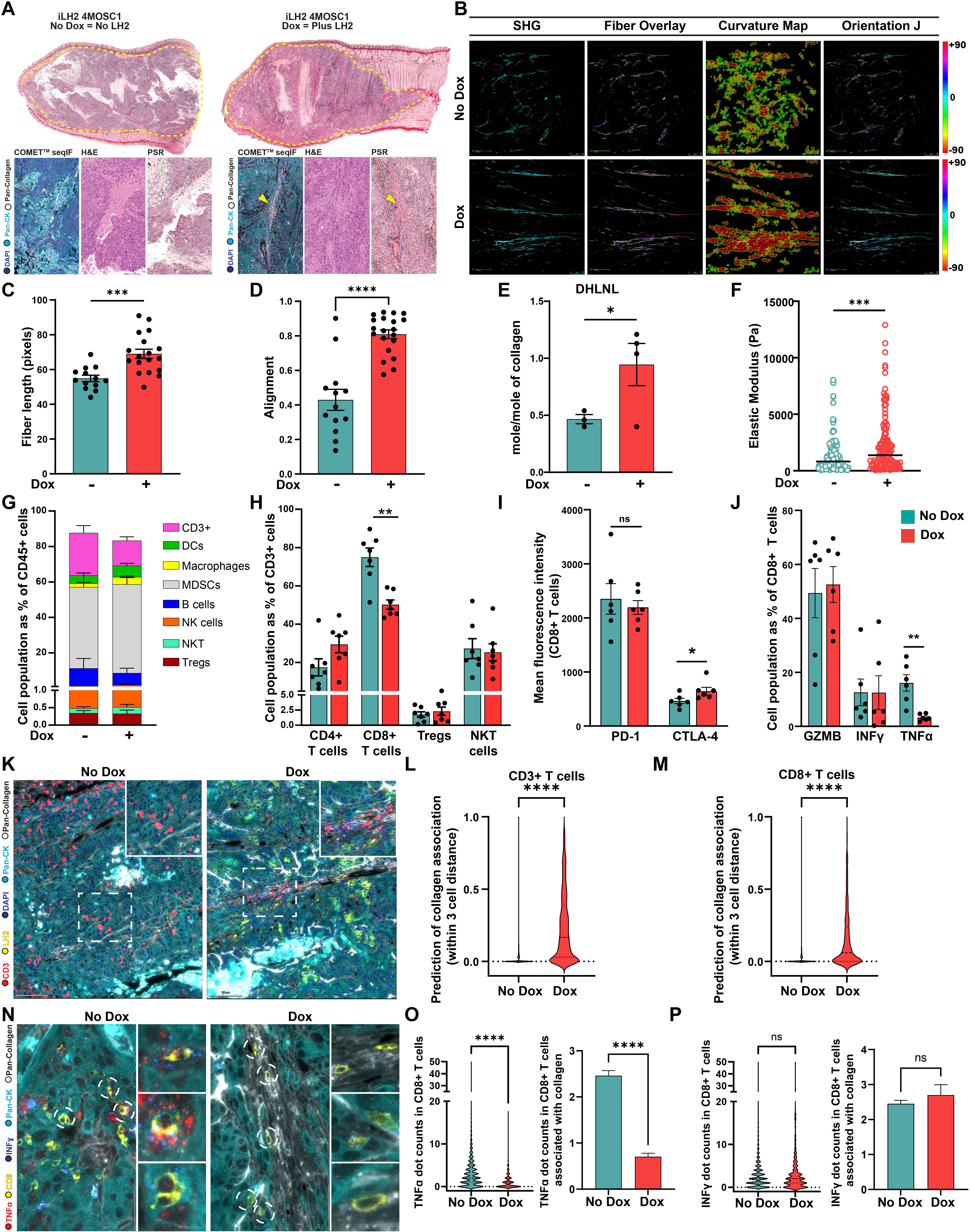
Inducible LH2 Expression in An Immunocompetent Murine Model Recapitulates HNSCC Patient ECM Remodeling and Immune Evasion. **(A)** *Top panel*, representative images of PSR stained sagittal sections of mouse tongue orthotopically transplanted with 4MOSC1 cells expressing inducible LH2, from mice maintained on regular chow (No Dox) or Doxycycline-containing chow (Dox). *Bottom panel*, ROIs depicting collagen architecture by multiplex IF imaging using Lunaphore COMET platform, H&E and PSR staining. **(B)** Representative SHG ROIs of mice tumors from No Dox and Dox conditions, with corresponding collagen fiber overlays (CT-FIRE), curvature maps (CurveAlign), and orientation-based color density overlays (OrientationJ). **(C)** Collagen fiber length analysis in mouse tumors under No Dox (n=5 mice) and Dox (n=5 mice) conditions, measured with CT-FIRE. Data points correspond to individual ROIs. Data points in the graph correspond to individual ROIs from the tumors. **(D)** Collagen fiber alignment analysis in mouse tumors under No Dox (n=5 mice) and Dox (n=5 mice) conditions, measured with CurveAlign. Data points in the graph correspond to individual ROIs from the tumors. **(E)** Mass spectrometry analysis of LH2-mediated DHLNL covalent cross-links in mice tumors from No Dox (n=3) and Dox (n=4) conditions. **(F)** AFM analysis of tumor stiffness in mouse tumors from No Dox (n = 3 mice) and Dox (n = 3 mice) conditions. Data point in the graph represents an individual indentation measurement. **(G)** Immune cell types represented as proportion of CD45+ cells infiltrating 4MOSC1 tumors from No Dox (n = 7 mice) and Dox (n = 7 mice) conditions. **(H)** Specific T-lymphocyte populations represented as proportion of CD3+ cells infiltrating 4MOSC1 tumors from No Dox (n = 7 mice) and Dox (n = 7 mice) conditions. **(I)** Mean fluorescence intensities of PD1 and CTLA-4 expressed on CD8+ T-cells infiltrating 4MOSC1 tumors from No Dox (n = 6 mice) and Dox (n = 6 mice) conditions. **(J)** Granzyme B, IFN-γ, and TNF-α expressing CD8⁺ T-cells are represented as a proportion of CD8⁺ T-cells infiltrating 4MOSC1 tumors under No Dox (n = 6 mice) and Dox (n = 6 mice) conditions. **(K)** Representative images from Lunaphore COMET multiplex staining of 4MOSC1 tumors (PCK+; cyan color) showing spatial distribution and proximity of T-cells (CD3+; red color) to collagen fibers (Pan-collagen+; white color) within the TME of control (No Dox) and LH2 (LH2+; yellow color) expressing Dox group. Nuclei are shown in blue. **(L-M)** Violin plots showing the predicted association of CD3+ and CD8+ cells with collagen within a 3-cell distance, as determined by Python-based analysis. **(N)** Representative images from Lunaphore COMET multiplex staining of 4MOSC1 tumors (PCK+; cyan color) showing spatial distribution and proximity of CD8+ T-cells (CD8+; yellow color) producing cytokines TNFα (*Tnf*α+, red color) and IFN-γ (*Ifng*+, blue) to collagen fibers (Pan-collagen+; white color) within the TME of control (No Dox) and LH2 (LH2+; yellow color) expressing Dox group. **(O)** Left, Violin plot showing abundance of *Tnf*α RNA transcript dots in CD8+ T-cells; Right, the predicted association of *Tnf*α*+* CD8+ cells with collagen within a 3-cell distance, as determined by Python-based analysis. **(P)** Left, Violin plot showing abundance of *Ifng* RNA transcript dots in CD8+ T-cells; Right, the predicted association of *Ifng*+ CD8+ cells with collagen within a 3-cell distance, as determined by Python-based analysis. Statistical analyses were performed using unpaired two-tailed Mann–Whitney U-test (C, D, H, I, J, L, M, O and P), unpaired one-tailed Student’s t test (E). Bar graphs represent mean ± SEM. ns, not significant. *P < 0.05, **P < 0.01, ***P < 0.001, ****P < 0.0001.

PSR staining showed no change in fibrillar collagen abundance, but polarized light imaging revealed a 1.6-fold increase in the proportion of mature collagen fibers (orange-red birefringence) in Dox-induced tumors compared to control (green-yellow) **(Figure S6A-C)**. SHG imaging revealed highly linear and aligned collagen fibers in the Dox group **(Figure 4B).** CT-Fire analysis showed increased fiber length **(Figure 4C),** alignment **(Figure 4D),** fiber number **(Figure S6D)** and reduced angular variation **(Figure S6E).** Curvalign analysis indicated higher kurtosis **(Figure S6F)** and aligned red pixels in the Dox group **(Fig S6G)**, confirmed by Orientation J **(Fig S6H)**. HPLC-based amino acid analysis indicates identical collagen content **(Figure S6I),** hydroxylysine (Hyl) levels **(Figure S6J)**, and aldehyde content **(Figure S6K)** between mice fed Dox chow versus those on standard chow (No Dox group). However, quantitative assessment of collagen cross-links revealed a significant elevation in LH2-mediated hydroxylysine aldehyde (Hyl^ald^)-derived cross-links (HLCCs), specifically DHLNL, within the tongue tumors of the Dox group compared to the No Dox group **(Figure 4E)**. A modest increase in the Hyl^ald^-derived trivalent cross-links, i.e. pyridinoline (Pyr) and deoxypyridinoline (d-Pyr), was observed in the Dox group, however, these differences did not reach statistical significance **(Figure S6L, S6M)**. Levels of HLNL, which can be derived from either lysine aldehyde (Lys^ald^) or Hyl^ald^, were comparable in both groups **(Figure S6N)**. The Lys^ald^ derived tetravalent cross-link (HHMD) did not change significantly between the two groups **(Figure S6O)**. Importantly, the ratio of HLCCs (DHLNL, Pyr, d-Pyr) to Lys^ald^-derived cross-links (LCCs: HHMD) was significantly elevated in the presence of LH2 overexpression **(Figure S6P).** A corresponding increase in the Hyl^ald^/Lys^ald^ ratio involved in cross-linking was also observed in the Dox group **(Figure S6Q)**. Atomic force microscopy (AFM) nanoindentation revealed a significant ∼2-fold increase in the elastic modulus of LH2-modified stroma in the tumors of Dox-fed mice **(Figure 4F).**

Flow cytometric evaluation reveals a diverse immune landscape comprising of cytotoxic T-cells (CD3+ CD8+), helper T-cells (CD3+ CD4+), B cells (CD19+ CD45R/B220+), regulatory T cells (Tregs-CD4+ CD25+ FOXP3+), natural killer cells (NK1.1+), NKT cells (NK1.1+ CD8+), DCs (CD11c+), macrophages (CD14+ F4/80+), as well as MDSCs (CD11b+ Ly6C^high^ Ly6G^high^) (**Figures 4G** and **S7A, S7B**), consistent with prior findings that 4MOSC1 tumors are highly immune-infiltrated (41). However, LH2-expressing tumors (Dox) exhibited significantly reduced CD8+ T-cell infiltration (7.87±3.56%) compared to control tumors (16.92±4.53%) in proportion to the total leukocyte (CD45+ cells) compartment **(Figure 4H)**. Immunofluorescence imaging of human HNSCCs, together with gene-expression correlations in the TCGA-HNSC dataset, corroborate this reduction **(Figure 3A, 3G)** suggesting that stiffened and aligned collagen in the LH2-high tumors may serve as physical barriers selectively limiting certain immune cells. Specifically, CD8+ T-cell migration within tumor parenchyma is limited such that cells are confined to the stromal compartment despite infiltration of other immune cell types similar to that observed in the human IE/F subgroup. Additionally, tumor-specific CD8+ T-cells in the Dox group exhibit modest increase in CTLA-4 expression, yet similar levels of PD-1 expression in both groups **(Figures 4I** and **S7C, S7D)**. Notably, a 5-fold decrease in the levels of TNF-α was observed in the CD8+ T-cells from the Dox group suggesting a dysfunctional phenotype mediated by alternative mechanisms of immune suppression **(Figure 4J)**.

High-plex spatial multiomics (RNA ISH + Protein IF) confirmed LH2 expression in the tumors (PanCK+ cells) of Dox chow fed cohort **(Figure 4K).** Pan-Collagen staining confirmed the abundance of collagen fibers in the TME and the effects of LH2 overexpression on their alignment. Interestingly, CD3+ T cells, specifically CD8+ T-cells, were observed to co-localize with the aligned collagen fibers in the Dox tumors, whereas the CD8+ T-cell infiltrates in No Dox tumors displayed no significant spatial correlation with collagen and were homogenously distributed throughout the tumor parenchyma **(Figure 4L, 4M).** Fluorescence *in situ* hybridization for detection of RNA transcripts confirmed a significant decrease in the *Tnf* counts in the CD8+ T-cells, especially in cells associated with aligned collagen fibers in the Dox group **(Figure 4N-O)**, while levels of *Ifng* remain unchanged **(Figure 4P)**. Interestingly, similar to observations made by flow cytometry **(Figures 4I** and **S7C, S7D)**, PD-1 and CTLA4 expression also did not differ much between the two groups as measured by seqIF staining suggesting an alternative mechanism is responsive for driving CD8+ T-cell dysfunction **(Figure S7E-G).** Collectively, this inducible syngeneic model recapitulates key features of human HNSCC including LH2 functionality, collagen architecture, and immune modulation making it a robust platform to investigate tumor-ECM-immune interactions.

### Exhausted ADGRG1+ CD8+ T-cells associate with LH2-modified fibrotic tissue contexts

Building on our previous analysis with ECM-high HNSCCs, the IE/F subtype was enriched for hypoxia, ECM-receptor interactions and PD-1 and PD-L1 checkpoint pathways, suggesting the operation of a fibrosis-driven mechanism of immune-suppression **(Figures 5A, B** and **S8A)**. Hypoxia regulates ECM remodeling (44), matrix deposition, immune responses (45), and transcriptional control of *PLOD2*/LH2 (21). Using the Hypoxia-Immune signature, we generated a Hypoxia score (46, 47) and observed that samples having high-Hypoxia score also had significant enrichment of the ECM score **(Figure S8B)**. DEG analysis stratified based on Hypoxia-Immune signature identified *PLOD2* and the adhesion GPCR gene *ADGRG1* as significantly upregulated in hypoxic HNSCC **(Figure 5C)**. Given the influence of GPCR signaling on cancer immunotherapy outcomes (48), we examined *ADGRG1* further and identified a positive correlation with *PLOD2* expression in the HNSC-TCGA **(Figure S8C)**. Further, *ADGRG1* is overexpressed in primary HNSCC tumors compared to adjacent normal tissue **(Figure 5D)** and this elevated expression correlates with poor overall survival **(Figure 5E).** In the Moffitt-IO cohort, patients treated with anti-PD1 monoclonal antibodies, namely Pembrolizumab or Nivolumab, showed significantly elevated expression of *ADGRG1* transcripts in the non-responders (n=27) compared to the responders (n=8), implicating a role for elevated LH2 and aberrant ADGRG1-mediated GPCR signaling in the failure of this form of immunotherapy **(Figure 5F)**.

**Figure 5.**
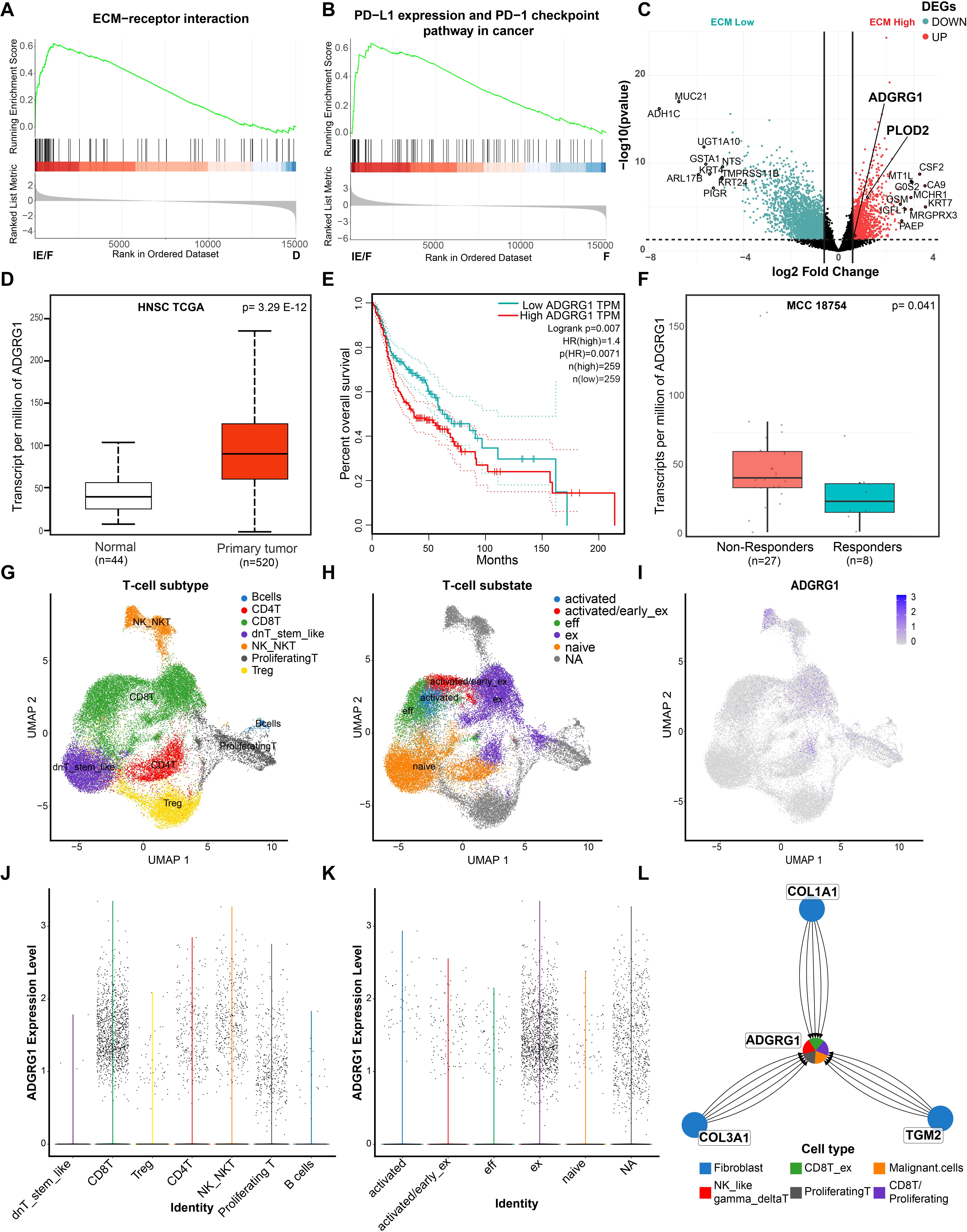
ADGRG1 expression in exhausted CD8+ T-cells in HNSCC corelates to poor survival outcomes and response to immunotherapy. **(A)** Pathway enrichment plot of ECM-receptor signaling in bulk RNA-Seq signature of IE/F vs F subtypes. **(B)** Pathway enrichment plot of PD-L1 expression and PD-1 checkpoint pathway in cancer in bulkRNA-Seq signature of IE/F vs F subtypes. **(C)** Volcano plot showing differentially expressed genes enriched in hypoxic ECM High tumor samples from Moffitt HNSCC TMA. **(D)** ADGRG1 expression in human matching normal tissue (n = 44; white) or primary tumor (n = 520; red) from TCGA Head and Neck Squamous Cell Carcinoma cohort (HNSC), Statistical analyses was performed using unpaired two-tailed Mann–Whitney U-test; ****P < 0.0001. **(E)** Kaplan-Meier depicting OS of ADGRG1 median stratified HNSC-TCGA patients (n=518). Univariate Cox Proportional Hazards Model hazard ratio (HR) and p value for ADGRG1 high vs low are indicated. **(F)** ADGRG1 expression in immunotherapy non-responders (n = 27; red) or responders (n = 8; cyan) from MCC18754 cohort, p value determined by two-sided Mann-Whitney test (****p<0.0001; Mann–Whitney Utest). Statistical analyses was performed using unpaired two-tailed Mann–Whitney U-test; *P < 0.05. **(G)** Uniform Manifold Approximation and Projection (UMAP) visualisation and clustering of 11,490 cells annotated as T-cell subtypes from integrated scRNA seq datasets. **(H)** UMAP clusters annotated as T-cell functional sub-states from integrated scRNA seq datasets **(I)** Feature plot showing the log-normalised expression of ADGRG1 over the UMAP structure and clusters confirming the source of ADGRG1 expression. **(G-H)** Violin plots showing ADGRG1 expression in T-cell clusters annotated as subtypes and sub-states, respectively. **(I)** Top ADGRG1 mediated receptor-ligand interactions identified by Cell-PhoneDB analysis in various cell types.

Interrogation of the Tumor Immune Single-cell Hub 2 (*TISCH2*) scRNA-seq database revealed *AGDRG1* expression within CD8+ T-cells and NK cell clusters from head and neck cancer datasets **(Figure S8D).** To delve further, we integrated human HNSCC scRNA-seq datasets using the Harmony package (v1.2.3) in Seurat (49, 50). A total of 36257 cells were jointly analyzed to annotate T-cell clusters according to gene expression profiles based on landmark genes **(Figure 5G).** This T-cell cluster was resolved into transcriptionally distinct sub-states **(Figure 5H)**, and *ADGRG1* is highly expressed in CD8+ T-cells **(Figure 5I, 5J)**, particularly within the exhausted CD8+ T cell subcluster **(Figure 5K)**. CellPhoneDB based cell-chat analysis identified fibroblast-derived (“sender”) *COL1A* (type I collagen) and *COL3A1* (type III collagen) as the top ligands for *ADGRG1*, with exhausted CD8+ T-cells as the “receiver” expressing this receptor **(Figure 5L).** Other notable ECM-receptor interactions included *FN* (fibronectin), *COL1A1*, *COL1A2*, *COL3A1, COL4A1*, *COL6A2*, *COL5A1* and *COL8A1* which can bind various integrin complexes **(Figure S8E, S8F).**

Tumors often hijack features of the wound healing process, including stromal remodeling, angiogenesis, and immune suppression and have been referred to as ‘wounds that do not heal’ (51, 52). To independently validate the presence of ADGRG1+ CD8+ T-cells in this context, a mouse wound healing dataset (GSE188432) was analyzed (53). CD3+ cells were sub-clustered and annotated based on curated gene signatures **(Figure S9A)**. *Adgrg1* transcripts were particularly enriched in the exhausted T-cell cluster **(Figure S9B, S9C)**. Interestingly, the fibroblast cluster (expressing *Col1a1*, *Fap*, *Fn1*) is associated with a progressive increase in *Plod2* expression going from day 4-7 in wounded samples compared to unwounded skin **(Figure S9D-H)**. These observations suggest that proper wound resolution depends on termination of T cell-mediated debridement and re-epithelialization that is controlled at later time points by LH2-driven collagen alignment and a mechanosensory immune checkpoint. Conversely, fibrotic disorders such as keloids can be viewed as ‘wounds that over-heal’, where excessive ECM deposition and dysregulated immune activity resembles features of the tumor stroma (54). Analysis of scRNA-seq datasets of human keloid patients and normal scar tissue (GSE175866 and GSE163973) (55) revealed elevated *PLOD2* levels in human keloid-associated fibroblasts as compared to normal scar tissue-associated fibroblasts **(Figure S9I-K)**, consistent with LH2-driven ECM stiffening that may prematurely dampen T-cell activity and impair tissue remodeling (56)

### A mechanosensory immune checkpoint is established by spatial localization of ADGRG1+ CD8+ T cells to LH2-modified collagen

To validate activation of ADGRG1 by type I collagen, we used a previously optimized serum response element (SRE)–luciferase assay (57) and confirmed that ADGRG1 is activated by type I collagen similar to its known agonists, type III collagen and P19 peptide, whereas the antagonist dihydromunduletone (DHM) blocks this activation (**Figure 6A**). Regulation of ADGRG1 by type I collagen was validated by spatial observations where CD8+ T-cells similarly associated with type I and type III collagen fibers in orthotopic 4MOSC1 tumors overexpressing LH2 (**Figure 6B**). Notably, type III collagen is co-expressed with type I collagen to form heterotypic bundles and regulates type I fibril diameter and architecture during fibrillogenesis (58, 59). To better understand the relevance of ADGRG1 in T-cell biology, we evaluated ADGRG1 expression in the tumor-infiltrating T-cell isolated from orthotopic models with or without LH2-modified collagen. Tumors from the Dox group with induced *PLOD2* overexpression had a >2-fold higher fraction of tumor-specific ADGRG1+ CD8+ T-cells as compared to the No Dox group **(Figure 6C, 6D**(*top panel*) and **S10A)**. A small subset of CD4+ T-cells and NK cells also expressed ADGRG1, however it did not differ between groups or across peripheral sites **(Figure S10A, S10B)**. ADGRG1+ CD8+ T-cells from the Dox tumors showed a significant decrease in the expression of markers of effector function, including TNFα (∼1.6 fold), IFNγ (∼2.3 fold), and Granzyme B (∼1.7 fold) as compared to cells from No Dox group **(Fig 6D, *bottom panel*)**.

**Figure 6.**
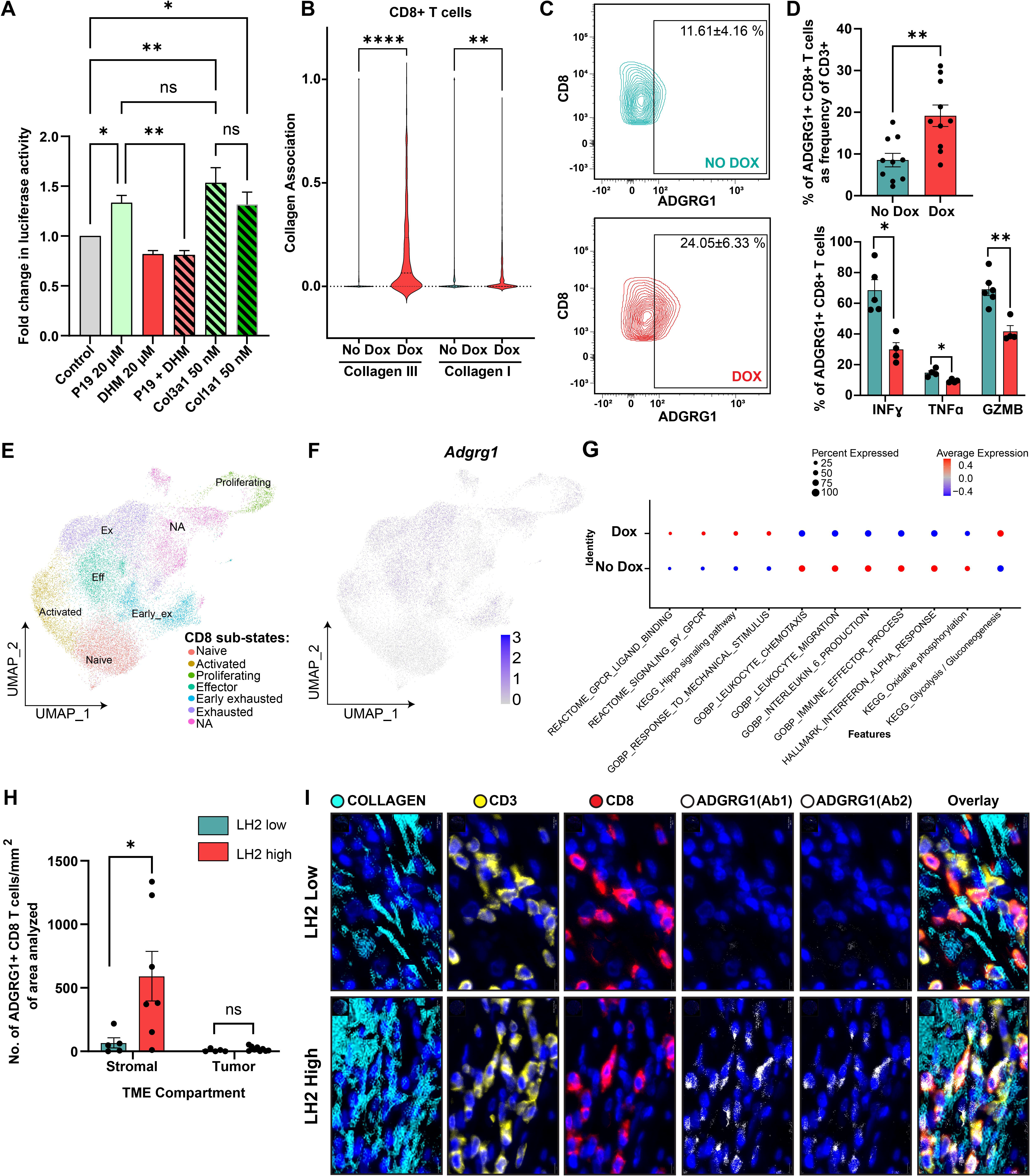
ADGRG1+ CD8+ T-cells spatially co-localize with aligned collagen fibers establishing a mechano-sensory inhibitory immune checkpoint. **(A)** HEK293T-cells transfected with plasmids to overexpress *Adgrg1* and an SRE-luciferase reporter were treated with DHM, P19, Col3a1 or Col1a1. P19 agonist-peptide activated SRE-luciferase but DHM inhibited the P19 stimulus. Error bars are mean ± SEM of three technical replicates. Statistical analysis was performed using One-way ANOVA; *P < 0.05, **P < 0.01. **(B)** Python based spatial distances were calculated and graphed as violin plots to compare spatial association of CD8+ T-cells with collagen I and collagen III in the No Dox and Dox samples. Statistical analysis was performed using One-way ANOVA; **P < 0.01, ****P < 0.0001. **(C)** Representative contour plots showing ADGRG1 expression in CD8+ T-cells infiltrating tumors of mice from the No Dox and Dox cohorts analyzed by flow cytometry. Percentages shown are Mean±SD values from No Dox (n=9) and Dox (n=7) mice. **(D)** ADGRG1 expressing CD8⁺ T-cells are represented as a proportion of CD3⁺ cells infiltrating 4MOSC1 tumors under No Dox (n = 6 mice) and Dox (n = 6 mice) conditions (top); IFN-γ, TNF-α and Granzyme B expressing ADGRG1+ CD8⁺ T-cells in tumors from No Dox (n = 6 mice) and Dox (n = 6 mice) cohorts (bottom). Statistical analysis was performed using two-tailed Mann-Whitney U test; *P < 0.05, **P < 0.01. **(E)** UMAP of unbiased clustering of 26930 CD8+ T-cells to identify clusters representing cellular sub-states distinguished by different colors. **(F)** UMAP showing expression of *Adgrg1* in CD8 T-cell sub states. **(G)** GSEA represented as dot plot displaying pathways enriched in exhausted T-cells from No Dox and Dox cohorts. **(H)** Bar graph showing absolute number of ADGRG1+ CD8+ T-cells in stromal and tumor compartments of LH2 low (n=5) and high (n=7) TMA cores. Statistical analysis was performed using two-tailed Mann-Whitney U test; *P < 0.05. **(I)** Representative images from Lunaphore COMET multiplex staining of TMA cores showing spatial distribution and proximity of ADGRG1+ CD8+ T-cells (CD3+: yellow; CD8+: red; ADGRG1: white) to collagen fibers (cyan color; overlaid using VALIS) within the TME of LH2 low and high cores.

To validate the effect of LH2-mediated ECM remodeling on T cell functionality, we performed scRNA-seq on tumors isolated from Dox and No Dox cohorts. *PLOD2* expression was confirmed in malignant cells (*Epcam*+ *GpNluc*+) from the Dox-induced cohort **(Figure S10C)**. Immune populations were annotated and visualized by UMAP **(Figure S10D)**. *Cd8a* expressing cells were sub-clustered into naïve, proliferating, activated, effector, early exhausted and terminally exhausted sub-states using curated exhaustion gene signatures **(Figures 6E** and **S10E, Table S9)**. Consistent with human HNSCC scRNA-seq data **(Figure 5L)**, *Adgrg1* expression is observed in exhausted, early exhausted, and effector CD8+ T-cell clusters in this mouse model **(Figure 6F** and **S10F)**. Quantification of *Adgrg1* expressing cells across CD8⁺ T-cell substates revealed low expression in naïve (8.2% in No Dox and 6.5% in Dox) and activated cells, with progressive upregulation towards more antigen-experienced states **(Figure S10G)**. Notably, CD8+ T-cells in the LH2-modified TME, particularly exhausted population, have the highest proportion of *Adgrg1* expressing cells (86.6% in Dox versus 53% in No Dox). Consistent with this murine tumor data, mIF based analysis of our human HNSCC TMA revealed a significant enrichment of ADGRG1⁺ CD8⁺ T-cells in LH2-high tumors (34%) as compared to LH2-low tumors (5.2%) **(Figure S10H)**.

GSEA revealed that exhausted T-cells are transcriptionally distinct between the two cohorts **(Figure 6G)**. Specifically, exhausted T-cells from LH2-expressing tumors (Dox) were significantly enriched for pathways associated with GPCR ligand binding and signaling, Hippo signaling and mechanosensitive responses and wound-healing–associated signatures **(Figures 6G, S9** and **S10I)**. Conversely, immune activation and effector function associated pathways like interleukin-6 (IL-6) production, interferon-α (INFα) response, and immune effector processes were downregulated in the Dox cohort **(Figure 6G)**. Notably, migration and chemotaxis-related genes were also reduced, consistent with physical confinement imposed by LH2-modified collagen matrices. Furthermore, depletion of oxidative phosphorylation and enrichment of glycolysis/gluconeogenesis indicates a metabolic shift toward glycolytic metabolism, often linked to T-cell dysfunction and adaptation to hypoxic or mechanically stressed environments (60, 61).

To confirm that ADGRG1 regulates the intra-tumoral distribution of CD8⁺ T-cells, spatial organization of ADGRG1+ cells was assessed in HNSCC patient samples by mIF. ADGRG1+ CD8+ T-cells predominantly localize to the stromal (PanCK⁻) compartment rather than to tumor regions **(Figure 6H)**, specifically colocalizing with aligned collagen fibers in the LH2-high tumors **(Figure 6I)**. Finally, TCGA breast and lung cancer datasets were analyzed as numerous studies demonstrate that both of these malignancies exhibit aligned collagen fibers, reflecting a common feature of tumor-associated matrix remodeling (62) (3, 63). *PLOD2* expression was markedly upregulated in the LUSC dataset compared to matched normal tissue, but remained unchanged in breast cancer (BRCA) samples suggesting that alternative modes of regulating LH2 expression and/or activity may be operational in BRCA (64, 65) **(Figure S11A)**. Notably, elevated *ADGRG1* expression correlated with poor overall survival in both cancer types **(Figure S11B, S11C)**. Moreover, *ADGRG1* was specifically upregulated in the CD8+ exhausted T-cell clusters in both malignancies, supporting broader PanCancer relevance of this mechanosensory checkpoint **(Figure S11D, S11E)**.

## Discussion

The TME is a dynamic and complex network of soluble factors and adhesion molecules that drive interactions between ECM components, stromal cells, tumor, and immune cells to augment cancer progression. The components of TME and their spatial organization collectively shape tumor architecture and progression. Although cancers have been extensively subtyped based on transcriptomic, epigenetic, mutational, and viral (HPV) profiles, (47, 66, 67) these classifications have largely overlooked the stromal landscape that forms the physical microenvironment of the tumor (68), particularly the ECM. To address this gap, we stratified HNSCC patients using an ECM-Immune signature revealing tumor phenotypes ranging from fibrotic immune-infiltrated to non-fibrotic immune-desert, each associated with distinct survival outcomes and response to immune-checkpoint blockade, highlighting the interplay between stromal architecture and immune infiltration in shaping the TME. Our approach is supported by a related PanCancer study by Bagaev et al., which defined four immune/fibrotic TME subtypes across 24 cancer types using transcriptomics data. The fibrotic subtypes appear to diverge functionally where the F subtype primarily establishes a physical barrier, with dense fibrillar collagen and additional ECM components (e.g., fibronectin, hyaluronan) that collectively exclude virtually all immune lineages, whereas in the IE/F subtype, the ECM acts as a contextual cue, constraining, guiding or functionally modulating infiltrating immune cells. Enrichment of LH2 within ECM-high phenotypes supports the view that LH2-mediated collagen crosslinking is a defining feature of fibrotic tumors, and within IE/F subtype, collagen organization directly reprograms CD8+ T-cell function rather than simply excluding these cells.

Within this context, we uncovered ADGRG1 as a key driver of immunosuppression in the fibrotic TME. Our findings align with PanCancer analysis by Wu et al. where 119 GPCRs were analyzed and *GPR56/ADGRG1* was observed to be amongst the GPCRs that strongly correlated with T-cell dysfunction score (48). Prior studies have described cell-intrinsic roles for ADGRG1, including suppression of NK cell effector functions (69), regulation of chemotaxis in cytotoxic and memory T-cells (70), and restriction of CD8⁺ T-cell migration (71). By integrating the stromal context, our findings extend beyond cell-intrinsic effects and highlight a role for ADGRG1 at the stromal–immune interface, linking receptor activity to ECM remodeling and CD8⁺ T-cell dysfunction, thereby explaining immune-suppressive mechanisms operational in IE/F subtype of HNSCCs that leads to poor responses observed to current ICIs. Interestingly, ADGRG1 upregulation was observed on tumor-reactive T-cells in acute myeloid leukemia (AML) suggesting an association with effector activity rather than immunosuppression (72). However, the absence of solid-tumor ECM constraints in AML limits its relevance to mechanisms operative in fibrotic cancers.

Our findings advance the understanding of TME heterogeneity and expand on the immunoregulatory role of ADGRG1 from a T-cell–focused checkpoint to a stromal–immune axis functioning as a mechanosensory checkpoint with therapeutic implications in fibrotic malignancies. This paradigm also resonates with non-malignant contexts such as wound healing and keloid formation, where excessive collagen cross-linking and alignment similarly suppresses CD8⁺ T-cell activity (73, 74). Specifically, key determinants of scar outcomes are linked to fibrillar type I collagen structural scaffolds that support immune-cell mediated debridement and re-epithelialization in early stages of wound healing, while later LH2 expression drives collagen alignment and wound resolution. In contrast, excessive LH2-mediated collagen cross-linking and alignment in keloids disrupt this progression, prematurely suppressing CD8⁺ T-cell function by imposing mechanosensory immune suppression similar to that observed in tumors **(Figure S12)**.

We acknowledge potential limitations in this study. Firstly, the association between TME-subtype and response to anti-PD-1 inhibitors is based on a retrospective dataset with a limited sample size. Secondly, although spatial profiling supports an anti-inflammatory microenvironment in LH2-modified tumors, the tongue-tumor bearing mice necessitate early euthanasia due to tumor burden, which likely prevented us from capturing later markers of systemic T-cell exhaustion such as INF-γ, even though reduction in TNFα levels was observed suggesting immunosuppression may already be underway at earlier time points (75). Finally, the it remains to be elucidated whether ADGRG1 is up-regulated in T cells in response to cues within the tumor microenvironment, or whether a pre-existing circulating ADGRG1⁺ T cell population becomes selectively enriched as it traffics through tumors containing LH2-modified collagen fibers. Further studies will be needed to distinguish between these possibilities.

Taken together, these findings position ADGRG1 as a promising therapeutic target that bridges immunosuppression and fibrotic pathologies, offering new opportunities for translational intervention. Since GPCRs are one of the most druggable protein families and already the focus of many clinical trials (76), our results provide a strong rationale for developing ADGRG1-targeted immunotherapies or repurposing existing GPCR-directed drugs to overcome immune evasion.

## Material and Methods

### HNSCC clinical samples and datasets

Clinical specimens were used from a previously published IRB-approved study of HPV-negative HNSCC patients at the H. Lee Moffitt Cancer Center(MCC#18754) (46).

Tissue microarray (TMA) of 121 cores (including 81 primary, 28 recurrent primary, and 12 lymph node metastases) was generated from pre-ICI biopsy punches; only primary tumors were analyzed to avoid radiotherapy-induced collagen changes in recurrent/metastatic cases. Matching bulk RNA sequencing data was available for 64 primary tumors from the Moffitt-IO cohort TMA; with additional samples from the Moffitt HNSCC cohort associated with the AVATAR dataset (n=125). Normalized RNA-sequencing data and clinical information for 515 TCGA-HNSCC cases were obtained from cBioPortal. All RNA-seq data were processed using a log2(TPM + 1) transformation.

### Hierarchical Clustering

ECM scores were generated using single sample gene set enrichment analysis (ssGSEA) with an ECM gene signature (**Table 1**) using the GSVA R package (77). Samples were stratified into ECM-high or ECM-low based on the median score. Primary tumors were classified into Immune-rich or depleted using TIMEx and Hallmark ssGSEA signatures. Expression values were z-score normalized across samples, clustered by unsupervised hierarchical clustering, and visualized with ComplexHeatmap.

### Gene Set Enrichment Analysis

GSEA was performed using ranked gene lists based on log2 fold-change values calculated with the DESeq2 R package (v1.42.1). We selected Hallmark signatures from msigdbr R package (v25.1.0). For KEGG signatures, the gene symbols were converted to entrez ID using the R package org.Hs.eg.db and clusterProfiler (v4.10.1). Enrichment analysis was conducted using clusterProfiler framework with a p-value cutoff of 0.05.

### Kaplan Meier curve

Overall Survival (OS) time was calculated in months from diagnosis to last follow-up or death, with follow-up beyond 60 months censored at 60 months. Survival difference was assessed using log rank test using ggsurvfit R package (v 1.1.0). GEPIA (Gene Expression Profiling Interactive Analysis) and UCSC Xena browser (https://xenabrowser.net/) were also used to plot gene specific OS of TCGA datasets. Only HPV negative samples were included.

### Picrosirius Red Staining

Slides were stained with Picrosirius Red (Polysciences, Cat# 24901-500) per manufacturer’s protocol. Briefly, sections were baked, deparaffinized using a Leica automated stainer, rehydrated, and stained with Weigert’s hematoxylin (8 min), followed by Solution-A (2 min), Solution-B (60 min), and Solution-C (2 min). After rinsing, slides were placed in 70% alcohol (45 secs), dehydrated and coverslipped. imaging was performed on a Zeiss Z2 Imager equipped with a polarizer, using an AxioCam 506 camera through a 20x/0.8NA objective lens. Tile scans generated whole slide images in polarized light and brightfield. Images were stitched with Zeiss Zen 2.3 and exported as TIFs for analysis with Image J CMM Lab PR-BRF Analyzer plugin (78).

### Second Harmonic Generation Imaging

H&E-stained sections were imaged on a Leica SP5 two-photon confocal microscope equipped with a MaiTai DeepSee Ti-sapphire laser tuned to 880 nm for SHG excitation, and emissions were collected by a non-descanned detector through a 440nm filter. Z-stacks were collected at 0.5 µm intervals, with a brightfield overlay captured with a 488 Argon laser and transmitted light PMT. Images were captured through a 25X/0.95NA water objective using Leica LASAF 2.6, and maximum projection of each z-stack were exported as TIF files. Collagen was quantified using CMM Lab-Collagen Fibril Orientation Analyzer (79), CT-Fire (v.2.0 beta) (https://loci.wisc.edu/software/ctfire) and CurveAlign (v5.0) (https://loci.wisc.edu/curvealign) modules.

### Multiplex Immunofluorescence using AKOYA

After the final strip, slides were counterstained with DAPI and coverslipped with ProLong Diamond. Imaging was performed on the Vectra®3 Automated Quantitative Pathology System. Formalin-fixed and paraffin-embedded (FFPE) samples were immunostained using the AKOYA OPAL 7-Color IHC kit on the BOND RX autostainer.

After baking at 65°C for 3h, TMA sections underwent automated deparaffinization, and antigen retrieval. For each marker, heat induced epitope retrieval (HIER) was performed (EDTA pH 9.0 buffer or citrate pH 6.0), followed by blocking and primary antibody incubation, LH2 (Proteintech, Rb Poly, 1:50, dye 570) at RT for 60 min followed by OPAL HRP polymer and one of the OPAL fluorophores during the final TSA step. Individual antibody complexes are stripped after each round of antigen detection. This was repeated five more times using the antibodies listed in **Supplementary Table S10**. After the final stripping, slides were counterstained with DAPI, and coverslipped with ProLong Diamond Antifade Mountant. Imaging was performed with the Vectra®3 Automated Quantitative Pathology Imaging System.

### Quantitative Image Analysis

Multi-layer TIFF from InForm (AKOYA) are imported into HALO for quantitative analysis. A classifier is trained to identify tumor, stroma or non-tissue regions, using Pan-cytokeratin as a tumor-masking marker. The tissue is segmented into individual cells using the DAPI marker which stains cell nuclei. For each marker, nuclear or cytoplasmic positivity thresholds are set based on expected staining patterns and published intensity ranges. Once thresholds are defined, the entire image set is analyzed with the created algorithm. The generated data includes positive cell counts per marker, percent of cells positive for the marker, and a per-cell analysis can be exported to provide the marker status, classification, and fluorescent intensities of every individual cell within an image.

### Virtual Alignment of Pathology Image Series (VALIS) and spatial statistics

Each sample included two serial TMA core images; one mIF stained and the other PSR stained. Images were registered using the VALIS pipeline (Figure 3D) (40) to align collagen fibers with tumor and stromal cells, then analyzed in QuPATH (80). Cells were segmented using the DAPI, object classifiers were created for each marker, and a pixel classifier was created to annotate collagen fibers. Centroid X-Y co-ordinates of CD3 and collagen annotations were used for spatial analysis.

Spatial statistic was performed in R (version 2025.05.0+596) using the “spatstat”. Gcross function quantified spatial proximity between CD3+ and PCK+ cells. Group level mean curves were plotted representing ±1 standard error compared to theoretical G(r) under complete spatial randomness. To evaluate the influence of the collagen, the ρ LJ (rou-hat) resource selection function was applied to assess how the density of CD3+ and PCK+ cells varies as a function of collagen intensity which is a spatial covariate in LH2 high and LH2 low tumors.

### Cell lines and cell culture

4MOSC1 mouse oral squamous cell carcinoma cells were kindly provided by Dr. Silvio J Gutkind at Moores Cancer Center, UC San Diego (41). Cells were cultured on Collagen I (Rat tail origin; Corning, 354236) coated plates in Keratinocyte Basal Medium 2 (PromoCell; C-20211) supplemented with the manufacturer provide SupplementMix and 0.06mM CaCl2 + 5ng/ml mouse recombinant Epithelial Growth factor (Thermo fisher, PMG8044) L+L1x Penicillin-Streptomycin-Glutamine (PSG; Thermo fisher, 10378-016)L+ 50 pM Cholera toxin (Millipore Sigma; C8052).

HEK 293T-cells were cultured in in Dulbecco’s modified Eagle’s medium (DMEM; GIBCO) supplemented with 10% FBS (Sigma). All cells were maintained at 37 °C in 5% CO2.

### Generation and validation of doxycycline inducible LH2 overexpression cells

4MOSC1 cells were initially transduced with a lentiviral carrying the EF1a-driven doxycycline-responsive reverse tetracycline transactivator 3 (rtTA3), following previously established protocols (28). Following puromycin selection, these cells were then spinfected with a retroviral doxycycline-inducible LH2 construct containing a constitutive GpNLuc LumiFluor cassette (see detailed protocol (28)). Fluorescence-activated cell sorting (FACS) was used to isolate GFP-positive, successfully transduced polyclonal population.

LH2 induction was confirmed at both mRNA and protein levels after 72 h of doxycycline treatment (1 mg/mL, replenished daily. Cells were harvested to confirm mRNA and protein expression. Unmodified ‘parental’ 4MOSC1 cells served as negative controls.

### qPCR

RNA was isolated using the NucleoSpin RNA kit (Macherey-Nagel, #740955) following the manufacturer’s protocol. RNA was eluted in RNase-free water and quantified using a Cytation 5 plate multimode reader spectrophotometer. cDNA was synthesized using the iScript cDNA Synthesis Kit (BioRad cat, #: 170-8891). Quantitative RT-PCR was performed using KAPA SYBR FAST qPCR Master Mix (KAPA Biosystems cat, #:KK4610), using 2 µL of cDNA (10 ng/µL) per reaction. qPCR primers used were as follows:

Human *PLOD2*.

Forward: GACCCGAGGAAGTGCTGAAG Reverse: CTTGTCGGCCAGTCTCTTGT

Mouse *Rpl23*.

Forward: GGC AAA CCA GAA CTA AGG AAA A Reverse: TTC GAT ATG ACT TTC GTT GTC G

### Western blotting

Cells were lysed in SDS Lysis Buffer [62.5 mM Tris-Cl, pH 6.8, 2% w/v SDS, 1% v/v glycerol] supplemented with protease (cat. #04693132001) and phosphatase inhibitors (cat. #10917400). Total protein concentration was quantified by BCA assay (Thermo Scientific, cat#; 23225). Approximately 20 µg of protein was resolved by SDS-PAGE. Membranes were incubated overnight at 4 °C with primary antibodies (anti-LH2, 1:100; β-actin, 1:10,000). HRP-conjugated secondary antibody (1:10,000) was applied for 1 h at room temperature. Protein bands were visualized using Clarity Western ECL Substrate (Bio-Rad, USA) and quantified with ImageLab 6.0.1 (Bio-Rad, USA).

### *In vivo* orthotopic xenografts

All procedures were approved by the University of South Florida Institutional Animal Care and Use Committee (IACUC) (11663R). Female C57BL/6 albino mice (8-9 weeks; Jackson Laboratories) were housed under specific pathogen–free conditions with water and food provided *ad libitum* at the H. Lee Moffitt Cancer Center. Mice were injected with 10^5^ accutase-dissociated single cells in 30 µL (15 µL PBS+L15 µL Matrigel) into the left lateral tongue under isofluorane anesthesia (2–2.5%). Animals in the Dox cohort were pre-fed Doxycycline chow (1000 ppm Doxy Diet, Envigo, cat# TD.180096) for 3 days prior to implantation and maintained on Dox chow. Animals were regularly monitored. Endpoints included >20% maximum body-weight loss or moribund condition. Tumor volume was assessed by calipers using ½ (length × width²). Lymph nodes and spleens were collected in 96-well plates for *ex vivo* analysis.

### In vivo bioluminescent imaging

Bioluminescent-fluorescent BRET signal was quantified noninvasively as previously described (28). Briefly, animals were i.p. injected with 250 μM (1:20 dilution, ∼500 μg/kg) Nano-Glo Luciferase Assay Substrate (Promega, cat. #N1120) in sterile DPBS. Isoflurane-anesthetized (3%) animals were imaged using the Newton 7.0 (Vilber) 5 min after injection. Mice were imaged using the following camera settings: Exposure time= 5 min, Binning= Medium, F-stop= 1.2, Field of view= 25. Data were analyzed using the Kuant software package. Signal was displayed as photons/s/cm^2^/sr.

### Atomic Force Microscopy

AFM measurements were performed using an Asylum Research Bio MFP-3D AFM integrated with a Nikon A1R inverted confocal microscope, mounted on an AVI-200 vibration-isolated and acoustically shielded platform. Cryopreserved mouse tumors were thawed, sectioned, and mounted onto glass slides using a thin layer of neutralized 2 mg/mL collagen I hydrogel, then incubated at 37°C for 20 minutes. The mounted samples were then placed on the temperature-controlled plate holder (Asylum Research), positioned on the inverted microscope stage, and maintained in warm culture medium (37°C) throughout the experiment. Silicon nitride probes (Novascan) with a nominal spring constant of 0.03 N m⁻¹ and a 4.5 µm polystyrene spherical tip were used, with spring constant calibrated by thermal tune method (81), and deflection sensitivity was determined in fluid against glass substrates serving as an infinitely stiff reference. Indentation was performed at 2.5 µm s⁻¹ with a maximum applied force of 3 nN. Each condition included at least 112 force–indentation curves, representing three technical replicates at each of a minimum of 37 randomly selected indentation sites. Elastic modulus was calculated using the Hertz model, assuming a Poisson’s ratio of 0.5.

### Collagen biochemical analysis

Tongue tumors were snap-frozen, pulverized in liquid nitrogen, washed and lyophilized. The dried samples were weighed, reduced with standardized NaB3H4, hydrolyzed with 6N HCl and subjected to amino acid and collagen cross-link analyses as described previously (28, 82–84). Collagen content and extent of lysine hydroxylation per collagen were determined by amino acid analysis. Reducible cross-links, dehydro-dihydroxylysinonorleucine (deH-DHLNL), deH-hydroxylysinonorleucine (deH-HLNL), and deH-histidinohydroxymerodesmosine (deH-HHMD) were analyzed as their radiolabeled, reduced forms, i.e. DHLNL, HLNL and HHMD. Non-reducible, stable cross-links, pyridinoline (Pyr) and deoxy-Pyr (d-Pyr) were simultaneously analyzed by their naturally occurring fluorescence. Total aldehydes, and HLCC-to-LCC ratios were determined following established protocols (26, 28).

### Tumor dissociation

Tumors were dissected and collected in Miltenyi MACS Tissue Storage Buffer (Miltenyi, Cat# 130-100-008). For preparation of single-cell suspensions, tumor tissue was manually minced using a scalpel, followed by enzymatic digestion with 1 mg/mL collagenase type I (Gibco, Cat# 17100-017) containing 5 mM CaCl₂ and 2.5 mg/mL dispase (Sigma, Cat# D4693) containing 50 mM HEPES-KOH (pH 7.4) and 150 mM NaCl. The solution was supplemented with DNase (2U/ml) and the ROCK inhibitor Y-27632 (1:1000). Minced tumor were transferred to a MACS C Tube (Miltenyi, Cat# 130-093-237) containing 5 mL of the collagenase/dispase solution and dissociation was performed on a gentleMACS Octo Dissociator (Miltenyi, Cat# 130-095-937) using a custom 12-min program at 37°C with alternating forward and reverse rotations (±200–400 rpm, 2 min per cycle). Enzymatic digestion was stopped by adding 5 mL of sterile 2 mM EDTA in PBS, and the cell slurry was transferred to a 15-mL conical tube and centrifuged at 750 × g for 5 min. The resulting pellet was resuspended in 5 mL Trypsin-EDTA (Sigma, Cat# T3924) and subjected to a second dissociation cycle on the gentleMACS using the same 12-min program. Trypsin was neutralized by adding 5 mL of complete medium, and the suspension was passed through 70-µm strainers to produce a single-cell suspension.

### Flow cytometry

Single-cell suspensions were washed with PBS and stained with Zombie viability stains (Biolegend, San Diego, CA). Cells were washed with cell staining buffer (Biolegend 420201) and resuspended in 90 µl in TruStain FcX CD16/32 (CLONE:93) and 10µl of Brilliant stain buffer (BD Biosciences 563794) for 15Lmin at 4L°C and protected from light. Cell surface staining was performed for 30Lmin at 4L°C with fluorophore-conjugated anti-mouse antibodies listed in **Supplementary Table S10**. Stained cells were washed and then fixed with BD cytofix (BD Biosciences, 555655) for 20Lmin at 4L°C, protected from light. In the case of intracellular staining, permeabilization was then performed by incubating with fixation-permeabilization buffer (BD Biosciences, cat#554715) according to the manufacturer’s recommendations prior to staining with intracellular targeted antibodies at the indicated dilutions in permeabilization buffer for 30Lmin at 4L°C and protected from light. Intracellular antibodies used are also mentioned in **Supplementary Table S10**. Cells were washed twice with permeabilization buffer and subsequently with cell staining buffer.

After a final wash, cells were resuspended in 500 µL of flow buffer and analyzed using Symphony II flow cytometer (BD Biosciences). Flow cytometry data were processed using FlowJo software (Tree Star). Cell populations were gated based on appropriate unstained and fluorescence-minus-one (FMO) controls.

### Integration of single cell datasets

Public sc-RNA-sequencing datasets of HNSCCs were downloaded from the Gene Expression Omnibus (GEO) (GSE181919 (Choi et. al), GSE103322 (85) and GSE164690 (86)).Cells with >10% mitochondrial gene content or <200 unique feature counts were removed using Seurat. The filtered dataset were combined and normalized using LogNormalize, (scale factor of 10,000), and subsequently log-transformed. Highly variable genes were identified, the data were scaled, and Principal Component Analysis was performed for dimensionality reduction. Batch effects were corrected with Harmony (v1.2.3) (50). Clustering was performed using Seurat’s shared nearest neighbor (SNN) method and visualized using UMAP plots. Cell typing was determined based on the differentially expressed genes of each cluster. Further, to examine *ADGRG1* expression across T-cell states, the major T-cell types previously annotated were sub-clustered into exhausted, naïve, activated or effector subsetss. The average expression of *ADGRG1* among these cells were visualized using violin plot.

### CellphoneDB

The ligand receptor interaction between various cell populations and CD8 exhausted cells was computed using CellphoneDB (v5) with a threshold of 0.1 and p-value of 0.05 (87). Statistical analysis method was implemented to infer significant cell-cell interactions, and the plots were visualized using the R package CCPlotR (v1.0.0). All the analyses were performed under R version 4.3.2.

### Luciferase Reporter Assays

HEK293T cells (1 × 10^5^) were seeded in a 12-well plate and co-transfected with *Adgrg1* plasmid (200ng; EX-Mm02830-Lv203, Genecopoeia), SRE-Luc reporter plasmid (100ng; pGL4.33[luc2P/SRE/Hygro] Vector, Promega) and mCherry plasmid (50ng; pSport6-mCherry, generated in-house) using Polyethylenimine (PEI); controls received-mCherry together with pcDNA3.1 empty vector. After 6hr of transfection, cells were serum-starved for 18hr and treated with DHM (antagonist), P19 peptide, Collagen III and Collagen I over the last 4 h of serum starvation. In the combination, cells were treated with P19 for 2 hrs and then DHM was added.

Cells were lysed in in 200 µL BrightGlo Luciferase Assay Buffer (Promega kit-E2650). From each lysate, 50 µL was transferred to an opaque white 96-well plate. mCherry fluorescence was measured first, followed by luciferase activity after adding 50 µL of BrightGlo Luciferase substrate (Promega kit E2650), using the Cytation 5 Hybrid Multi-Mode Reader (BioTek). Luciferase values were normalized to mCherry and expressed relative to no-reporter controls.

### Single-cell cDNA and library preparation and sequencing

Single-cell RNA sequencing was performed using the Chromium System (10x Genomics, Pleasanton, CA) at the Molecular Genomics Core, Moffitt Cancer Center and Research Institute. Following tissue harvest and dissociation, single-cell suspensions were prepared, washed twice with 1× PBS (calcium- and magnesium-free) containing 0.04% (w/v) BSA, and resuspended in the same buffer according to the 10x Genomics cell preparation protocol. Cell viability and concentration were determined using acridine orange/propidium iodide (AO/PI) dual fluorescent staining on a Cellometer K2 (Nexcelom Bioscience LLC, Lawrence, MA). Cells were then loaded onto the Chromium Single Cell Controller at a concentration of 1,000 cells/µL, targeting a recovery of approximately 10,000 cells per sample.

Individual cells, reagents, and gel beads were encapsulated into nanoliter-scale Gel Beads-in-Emulsion (GEMs), and reverse transcription of polyadenylated mRNA was performed within each droplet at 48 °C. Gene expression (GEX) libraries were prepared following the Chromium GEM-X Single Cell 5′ v3 Gene Expression User Guide. Sequencing was conducted on an Illumina NovaSeq 6000 platform, generating approximately 50,000 reads per cell.

### Single-cell RNA-sequence data processing

Data processing—including demultiplexing, barcode assignment, alignment, and gene counting—was performed using Cell Ranger. Reads were mapped to a custom mouse reference genome (GRCm39 with human *PLOD2* sequence added and mouse *Plod2*removed). Feature-barcode matrices were generated with Cell Ranger; visualized and analyzed in Loupe Browser v9.0. Cells with <200 or >9000 features or >20% mitochondrial content were removed. Subsequent steps followed the Seurat workflow as mentioned above. The immune cells were identified by *Ptprc* expression;the cells were subclustered and annotated based on the DEG list using canonical markers for Monocytes/Macrophages (*Cd68* and *Cd14)*, Dendritic cells (*Itgax*), NK cells *(Ncr1* and *Ncam1)*, and T-cells (Cd3g and Cd8a). The cell states were annotated using module scores of certain curated exhaustion markers using Seurat’s AddModuleScore function.

Difference in the exhausted T-cells between Dox and No-Dox conditions were assessed by comparing module score derived from immune signatures in MsigDB and KEGG.

### Lunaphore COMET multiplex IF

FFPE slide was heat fixed at 60°C overnight on ACD HybEZ II Hybridization System (Bio-Techne). Heat-fixed slide was processed with PT Module (Epredia) using Dewax and HIER Buffer H (TA999-DHBH, Epredia) for 60 min at 99°C, rinsed, and stored in Multistaining Buffer (BU06, Lunaphore). The multiplex RNA+protein template was generated using the COMET Control Software, and reagents were loaded to perform fully automated RNAscope^TM^ HiPlex Pro Assay (Advanced Cell Diagnostics, Inc.) and sequential immunofluorescence (seqIF) protocol (88, 89). Samples were pretreated in the RNAscope™ HiPlex PretreatPro buffer for 30 minutes at 40°C, followed by hybridization of mixed RNAscope™ HiPlex target probes to *Mus musculus* (*Ifng-*T4, *Tnfa-*T1) for 2 h at 40°C. Detection used HiPlex Pro Fluoro 4-channel probes for 15 min at 37°C, washing in 1× MSB at 27°C, and imaging [FITC, TRITC, Cy5, Cy7]. Fluorophore cleavage was performed with HiPlex Pro Cleaving Reagent (1:10 in 4× Saline Sodium Citrate) for 15 min at 27°C, followed by washing; detection/cleavage cycles were repeated three times. After final RNA imaging and tissue-background imaging, the seqIF™ protocol began.

Nuclear signal was detected with DAPI (Thermo Scientific, cat no: 62248, 1/200 dilution). For all staining cycles, the dynamic incubation time of primary antibody mixes was set to 4 min, while the dynamic incubation time of secondary antibodies and DAPI cocktails was set to 2 min. All primary antibody cocktails were diluted in Multistaining Buffer (BU06, Lunaphore). For each imaging cycle, the following exposure times were used: DAPI 80 ms, TRITC 400 ms, Cy5 200 ms. The elution step duration was set to 2 min for each cycle and was performed with Elution Buffer (BU07-L, Lunaphore). The quenching step was set to 30 s and was performed with Quenching Buffer (BU08-L, Lunaphore). The imaging step was performed with Imaging Buffer (BU09, Lunaphore). ntibodies are listed in **Supplementary TableS10**. A raw OME-TIFF file was generated by COMET Control software for downstream analysis.

### Image pre-processing and analysis

The final COMET workflow includes alignment, stitching, flat-field correction, and generation of output 16-bit OME-TIFF images in the COMET Control software after the automated RNAscopeTM HiPlex + seqIF protocol execution and data acquisition. Pixel-wise autofluorescence correction for each marker was performed using Horizon Viewer. Autofluorescence images acquired before each imaging cycle were used for correction to minimize the occurrence of background subtraction artifacts. This also allowed us to ensure the minimal deviation of the fluorescence intensity values that might be caused by the and reduce photobleaching related intensity drift.

Cell segmentation was conducted in Horizon or QuPath, following previously established workflow for detection and classification of individual cells. The spatial positions (X–Y coordinates) of segmented cell were exported and analyzed in Python using custom scripts (90–92) to assess spatial organization, cell–cell proximity, and neighborhood composition within the TME. In brief tables were used in pandas format for easy visualization, basic calculations (Average radius/Averages etc.) were determined using the Numpy package and cell proximity predictions were determined using radius_neighbors. A value of 0 to 1 was given to each cell depending on the proximity to the target cell type to a given radius distance. An average cell radius per sample was used as the reference scale for all proximity calculations.

### Statistical Analysis

GraphPad Prism was used for statistical analyses. Differences between groups were evaluated by unpaired 2-tailed Student’s t test or one-way ANOVA for repeated measures. Data are presented as the mean ± SEM. P values of less than 0.05 were considered statistically significant.

## Data Availability

The raw data for scRNA-seq reported in this article have been deposited in the Gene Expression Omnibus repository (Accession Number:xxxx). All other data and materials are available from the corresponding author.

## Authors’ Disclosures

S. Srivastava and A.L. Amelio declare a patent application related to this work. M.C., P.B. and J.G. are employees of Bio-Techne, which manufactures reagents and the COMET instrument used in this study, and MC and PB are also shareholders. C.H. Chung and R.J.C. Slebos report honoraria from Fulgent, Genmab, AVEO, Seagen, Regeneron, Bicara, Johnson and Johnson, and Exelixis for Scientific Advisory Board participation. The remaining authors declare that they have no conflict of interest.

## Acknowledgements

The authors gratefully acknowledge the patients who participated in this study, without whom this work would not have been possible. We also wish to thank Jodi Kroger, Bethany Carter, and Neelkamal Chaudhary of the Flow Cytometry Core, Jodi Balasi of the Tissue Core, Carlos M. Moran Segura, Jonathan V Nguyen, Neale Lopez-Blanco and Anthony Alleyne of the Advanced Analytical and Digital Pathology, Sean J Yoder, Chaomei H. Zhang, Lan M. Zhang, and Tania E. Mesa of the Genomics Core, Mahmoud Abdalah of the Quantitative Imaging Core, and Mikalai Budzevich and Epi Ruiz of the Small Animal Imaging Lab at the H. Lee Moffitt Cancer Center and Research Institute for their expert technical assistance. We also thank Drs. Ignacio Wistuba, Alex Jaeger, and Vince Luca, and members of the Amelio Lab for helpful discussions, suggestions, and/or scientific review of this article. The RNA-seq data included in this study was obtained through the Oncology Research Information Exchange Network (ORIEN) Avatar Project initiated under the Total Cancer Care protocol at the Moffitt Cancer Center. This work was supported in part by a Florida Biomedical Research Program James & Esther King grant (21K04) (C.H.C.), the Moffitt Cancer Center’s “Center of Excellence for Evolutionary Therapy” (J.W.), the Moffitt Cancer Center’s “Cancer Biology & Evolution Program” pilot award (J.W. and A.L.A.), and Moffitt Cancer Center startup funds (to A.L.A.).

## Author Contributions

Conception and design, S.S, and A.L.A.; development of methodology, S.S., J.J.P., S.M., R.N., H.B., F.F., M.Y., J.W., J.S.G., and A.L.A.; acquisition of data (provided reagents, provided facilities, etc.), S.S., J.J.P., R.N., L.S.K., J.T., R.C., M.C., D.T.N., M.Y., J.W., J.S.G., C.H.C., A.L.A.; interpretation of data (e.g., statistical analysis, biostatistics, and computational analysis), S.S., J.J.P., J.H.M., S.M., X.S., J.A.G-P., D.T.N., M.Y., J.W., J.S.G., C.H.C., A.L.A.; writing of the manuscript, S.S. and A.L.A.; review and revision of the manuscript, S.S., J.J.P., J.H.M., S.M., R.N., L.S.K., J.T., H.S.B., F.F., R.C., P.B., J.G., R.J.C.S., J.O.J., C.G., X.S., M.C., J.A.G-P., D.T.N., M.Y., J.W., J.S.G., C.H.C., A.L.A.; administrative, technical, or material support (i.e., reporting or organizing data and constructing databases), S.S., J.J.P., J.H.M., P.B., J.G., M.C.; study supervision, S.S. and A.L.A.; acquisition of funding, A.L.A.

---

**Supplementary Figure 1. TME subtype identification in Moffitt-HNSCC and TCGA-HNSC cohorts**. Heatmaps representing TME subtypes in the **(A)** Moffitt-HNSCC patient cohort (n = 125) and **(B)** the TCGA-HNSC cohort (n = 515). RNA expression data were z-normalized, each row represents a single gene signature from TIMEx and MSigDB Hallmark Pathways, each column represents a patient sample. **(C)** Kaplan–Meier survival plot for OS of HNSC-TCGA patient cohort segregated based on median of ECM score into high (fibrotic) and low (non-fibrotic). **(D)** Kaplan–Meier survival plots for OS of HNSC-TCGA patients classified as IE TME subtype compared to F subtype, **(E)** IE/F subtype and **(F)** D subtype. P values were calculated by Log-rank (Mantel-Cox) test.

**Supplementary Figure 2. GSEA comparison of TME subtypes in Moffitt and TCGA cohorts.** Hallmark and KEGG pathways enriched in **(A-B)** Fibrotic versus Immune depleted subtype, **(C-D)** Fibrotic - Immune enriched vs Immune enriched subtype and **(E-F)** Fibrotic - Immune enriched vs Fibrotic in HNSC TCGA and Moffitt-HNSCC cohorts; NES represents the normalized enrichment score, as computed by GSEA; p value <0.05.

**Supplementary Figure 3. *PLOD2* is a key driver of collagen architecture in fibrotic HNSCCs. (A)** Volcano plots of ECM-associated DEGs enriched in ECM-high tumors across TCGA-HNSC and **(B)** Moffitt-HNSCC cohorts. **(C)** Schematic illustrating LH2-mediated collagen crosslinking in HNSCC, created using BioRender.

**Supplementary Figure 4. LH2 mediated collagen alignment influences immune cell distribution in HNSCC TME. (A)** Representative images of Multiplex immunofluorescence (IF) staining of CD14+ (yellow), CD11c+ (green) and Pan-CK (cyan) cells in HNSCC tumor cores stratified by LH2 expression. **(B)** Distribution of CD14+ cells and **(C)** CD11c+ cells across tumor (PCK+) and stromal areas (PCK-) of LH2 low and LH2 high cores **(D)** Gcross analysis of CD3+ to PCK+ cell proximity across LH2 groups; dashed line denotes spatial independence. **(E)** Functional ANOVA of G(r) curves across LH2 groups, showing -log10(p-value) within distances. Red points mark distances where T cell-tumor cell association differs significantly between groups **(F)** Resource selection analysis using rho-hat function showing the relative intensity of PCK+ cells as a function of local collagen density (within one cell diameter), which appears similar between LH2-high and LH2-low tumors. **(G)** Correlation between PLOD2 expression and infiltration of CD8+ T-cell, **(H)** CD4+ T-cell, **(I)** NKT-cell, **(J)** monocytes/macrophages, **(K)** myeloid-derived suppressor cells (MDSCs), and **(L)** dendritic cells using deconvoluted TCGA HNSCC data. Statistical analyses was performed using unpaired two-tailed Mann–Whitney U-test (B-C). Error bars are mean ± SEM; ns, not significant, **P < 0.01. TIMER plots indicate computed Spearman’s rho and p-values.

**Supplementary Figure 5. LH2 overexpression in murine 4MOSC1 cells to establish an immunocompetent orthotopic *in vivo* model. (A)** Quantitative real-time PCR analysis of *PLOD2* mRNA levels in unmodified (parental) and LH2-overexpressing 4MOSC1 cells treated +/- 1 μg/mL doxycycline for 72h. *PLOD2* expression was normalized to *Rpl23* mRNA levels and fold expression was calculated relative to the parental cells. **(B)** Total protein extracted, and western blot run to validate doxycycline induced LH2 expression at the protein level in 4MOSC1-*PLOD2* cells compared to parental cells. This blot is representative of independent biological replicates**. (C)** Quantification of LH2 protein levels. Band intensities were normalized to β-actin and are shown as the fold change relative to the no doxycycline condition in parental cells. **(D)** Representative *in vivo* bioluminescent images (BLI) of GpNluc-labeled 4MOSC1 cells at various times post-orthotopic tongue xenograft transplantation in animals maintained either on normal or doxycycline chow. **(E)** BLI signals represented as fold change for each imaged time point. **(F)** Endpoint tumor volume defined as 1/2(length ×width^2^) was determined by external caliper., **(G)** Representative *ex vivo* BLI images of cervical lymph nodes resected from control (No Dox) and doxycycline-chow (Dox) fed animals at study endpoint **(H)** Quantification of metastatic lymph node *ex vivo* BLI signals by ROI analysis of images obtained at endpoint (n = 5 mice per group). Statistical analyses was performed using One-way ANOVA test (A), unpaired two-tailed Mann–Whitney U-test (E, F, H). Error bars are mean ± SEM; ns, not significant, **P < 0.01, ****P < 0.0001.

**Supplementary Figure 6. Syngeneic mice model recapitulates collagen architecture of human HNSCC. (A)** Representative ROIs of 4MOSC1 tumors from animals administered either normal (n=5) or doxycycline chow (n=5) stained with Picrosirius Red (PSR). **(B)** Total collagen area was quantified using images captured under bright-field and **(C)** collagen birefringence was quantified based on red or green signals generated under polarized light microscopy. **(D-E)** CT-FIRE analysis compared individual fiber numbers and angular variance, **(F–G)** CurveAlign module compared collagen fiber kurtosis and pixel intensity, and **(H)** OrientationJ measured the normalized fibril orientation distribution between No Dox and Dox groups. **(I-K)** Total collagen cross-link quantities were compared in 4MOSC1 tumors from animals administered either normal (n=3) or doxycycline chow (n=4). Quantification of total collagen cross-links, the ratio of hydroxylysine (Hyl) to hydroxyproline content (Hyp/Hyp x 300) and total aldehyde levels. **(L-P)** Analysis of collagen cross-link quality in 4MOSC1 tumors from animals administered either normal or doxycycline chow. Quantification of Pyr, d-Pyr, HLNL, HHMD and the ratios of HLCC-to-LCC [(DHLNL + Pyr + d-Pyr)/HHMD] and ratio of Hyl^ald^/Lys^ald^ [(DHLNL + Pyr x 2 + d-Pyr x 2)/HHMDx2] were calculated. Statistical analyses were performed using unpaired two-tailed Mann–Whitney U-test (B-G) and One-tailed unpaired t-test (I-Q). Bar graphs are mean ± SEM; ns, not significant, *P < 0.05, ****P < 0.0001.

**Supplementary Figure 7. LH2-mediated collagen alignment hinders infiltration of tumor specific CD8+ T-cells and drives their exhaustion in syngeneic mouse model. (A)** Immune cell types represented as proportion of CD45+ cells from Lymph node, and **(B)** spleen of tumor bearing animals from No Dox (n = 7 mice) and Dox (n = 7 mice) groups **(C)** Mean fluorescence intensities of PD1 and CTLA-4 expressed on CD8+ T-cells Lymph nodes, and **(D)** spleen of tumor bearing animals from No Dox (n = 6 mice) and Dox (n = 6 mice) conditions. **(E)** Representative images from COMET multiple IF imaging showing CD8 (yellow), PD1 (red), CTLA4 (blue), Pan-collagen (white) and Pan-CK (cyan) **(F)** Spatial correlation of collagen with cd8+ T-cells expressing PD1 and, **(G)** CTLA4. Statistical analyses was performed using unpaired two-tailed Mann–Whitney U-test. Bar graphs are mean ± SEM; ns, not significant, **P < 0.01.

**Supplementary Figure 8. ADGRG1+ exhausted CD8+ T-cells interact with collagen in the TME of HNSCC. (A)** Pathway enrichment plot of hallmark_Hypoxia signaling in bulkRNA-Seq signature of IE/F vs F subtypes. **(B)** Box plot comparing enrichment of hypoxia score in bulk RNA seq data from ECM high and low samples. **(C)** Correlation plot for log 2 normalized expression of *PLOD2* and *ADGRG1* transcripts in TCGA-HNSC dataset. **(D)** *ADGRG1* expression in HNSC & NPC scRNA seq datasets available on TISCH2. **(E)** Receptor-ligand interactions identified by Cell-PhoneDB analysis in CD8 exhausted as receiver. **(F)** Top overall receptor ligand interactions. Statistical analyses was performed using unpaired two-tailed Mann–Whitney U-test **(B)**. Error bars are mean ± SEM; **P < 0.01. Correlation plots indicate Spearman’s rho and p-values computed by GEPIA **(C)**.

**Supplementary Figure 9. LH2-driven matrix remodeling and immune suppression are recapitulated in wound healing. (A)** Integrated UMAP representing CD3 cell clusters from all wound timepoints in GSE188432 annotated as T-cell sub-states. **(B)** Feature plot showing *Adgrg1* expression in T-cell sub-state clusters. **(C)** Dot plot showing expression of genes or gene signatures used for annotation of T-cell sub-state clusters. **(D)** Integrated UMAP representing all cells from unwounded skin and skin collected 4 days or 7 days post-wound. **(E)** Feature plots of cells from all wound healing time points showing expression of *Fap*, *Col1a1* and *Fn1* for identification and annotation of fibroblast cluster. **(F)** Feature plot showing *Plod2* expression in unwounded samples and samples collected **(G)** 4 days post wound and **(H)** 7 days post wound. **(I)** Integrated UMAP for keloid (n=3) and normal scar (n=3) samples from GSE163973 with original annotations (inset). **(J)** Feature plot showing PLOD2 expression in normal scar fibroblasts (NF1-3) and **(K)** keloid fibroblasts (KF1-3).

**Supplementary Figure 10. Tumor-specific ADGRG1+ exhausted CD8+ T-cells are enriched in LH2 modified TME. (A)** ADGRG1 expression in CD8+ T-cells, CD4+ T-cells, NKT-cells and NK cells are represented as a proportion of CD45⁺ cells infiltrating tumors of mice from the No Dox (n=11) and Dox (n=11) cohorts analyzed by flow cytometry. Statistical analysis was performed using two-sided Mann-Whitney test, **P < 0.001. **(B)** Frequency of ADGRG1+ CD8+ in blood, spleen and lymph nodes harvested from healthy mice and tumor bearing mice from no dox and dox cohorts represented as a proportion of CD45⁺ cells. **(C)** *PLOD2* expression in malignant cells (GpNluc+) from No Dox (n=3) and Dox (n=3) cohorts. **(D)** UMAP showing all *Ptprc+* (CD45+) cells from No Dox (n=3) and Dox (n=3) cohorts annotated as various immune cell types based on expression of canonical markers in the cell cluster. The number in the parenthesis indicates the cell counts from No Dox (cyan) and Dox (red) cohorts. **(E)** Dot plot showing expression of genes or gene signatures used for the annotation of CD8+ T-cell clusters into sub-states. **(F)** Feature plot showing *Adgrg1* expression in all CD45 cell clusters from No Dox (n=3) and Dox (n=3) cohorts. **(G)** Heat map showing *Adgrg1* expressing cells in the respective sub-state as proportion of total cells in the cellular sub-state cluster. White represents the lowest value, and Red represents the highest value. **(H)** Frequency of ADGRG1+ CD8+ cells represented as proportion of total CD8+ T-cells in LH2 low (n=5) and high (n=7) TMA cores. **(I)** Dot plot showing enrichment of vascular wound healing and atypical scaring of skin gene signatures in exhausted T-cells from Dox cohort. Statistical analysis was performed using two-sided Mann-Whitney test (A, H) and One-way ANOVA (B). ns, not significant; **P < 0.01.

**Supplementary Figure 11. Collagen stiffness–ADGRG1 axis drives CD8**⁺ **T-cell exhaustion across cancers. (A)** Expression of *PLOD2* in TCGA breast and lung cancer samples compared to matched normal tissue. **(B)** Kaplan-Meier depicting OS of *ADGRG1* median stratified TCGA patients from BRCA and **(C)** LUSC. Univariate Cox Proportional Hazards Model hazard ratio (HR) and p value for *ADGRG1* high vs low are indicated. **(D)** *ADGRG1* expression in sc-RNA seq datasets of breast cancer and **(E)** Lung squamous cell carcinoma, available on TISCH2.

**Supplementary Figure 12. Model of ADGRG1 in LH2-modified fibrotic tissue contexts.** Illustration highlighting similarities between the proposed mechanosensory immune checkpoint linked to stromal remodeling and immunosuppression of ADGRG1+ CD8+ T-cells in oral carcinogenesis and wound healing contexts, created using BioRender.

