## Supplementary figures and images for "Collagen remodeling promotes a GPCR-mediated mechanosensory immune checkpoint in ADGRG1+ CD8+ T cells and serves as a spatial biomarker of response to immunotherapy"

### Supplemental Figure S1

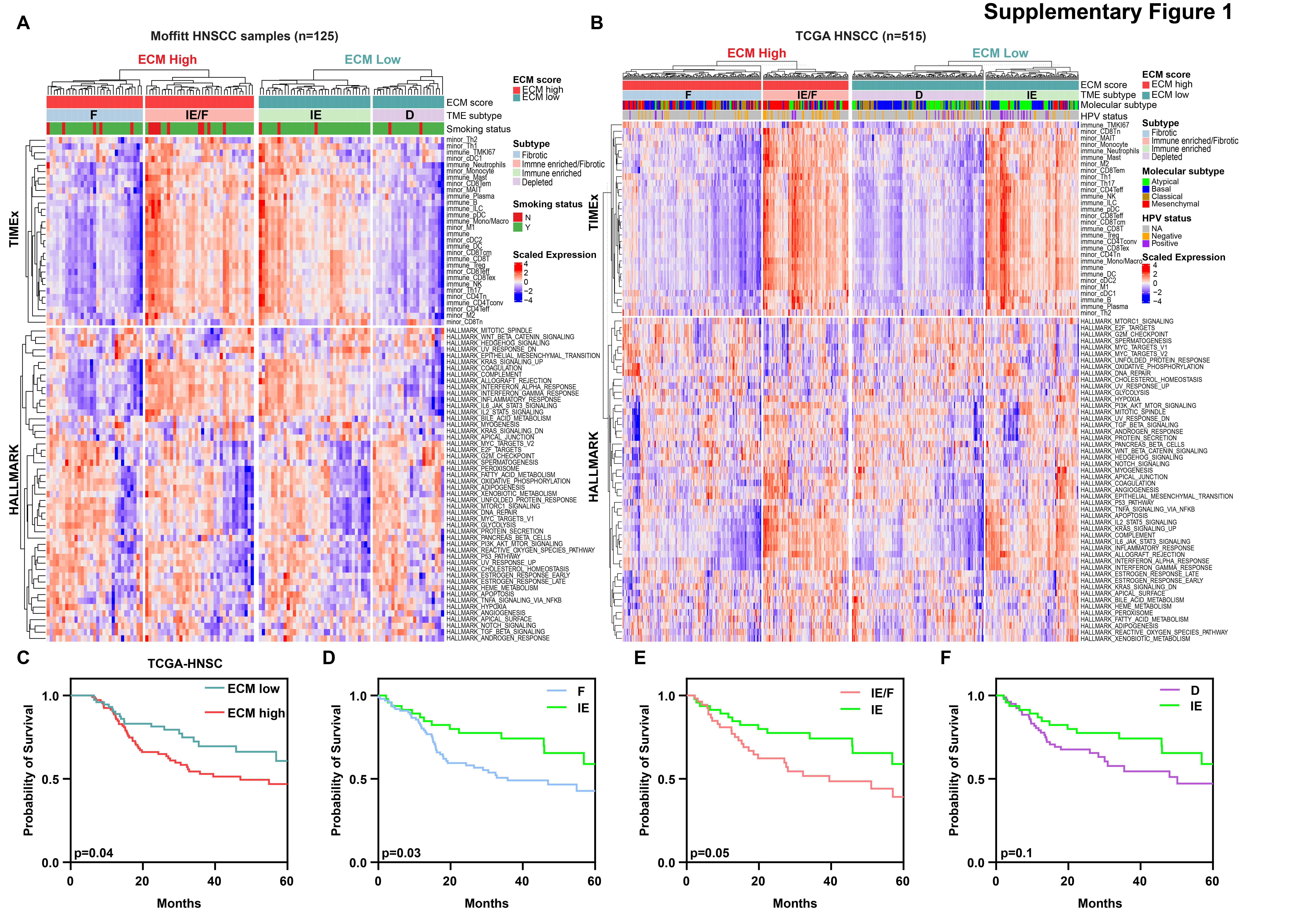

### Supplemental Figure S2

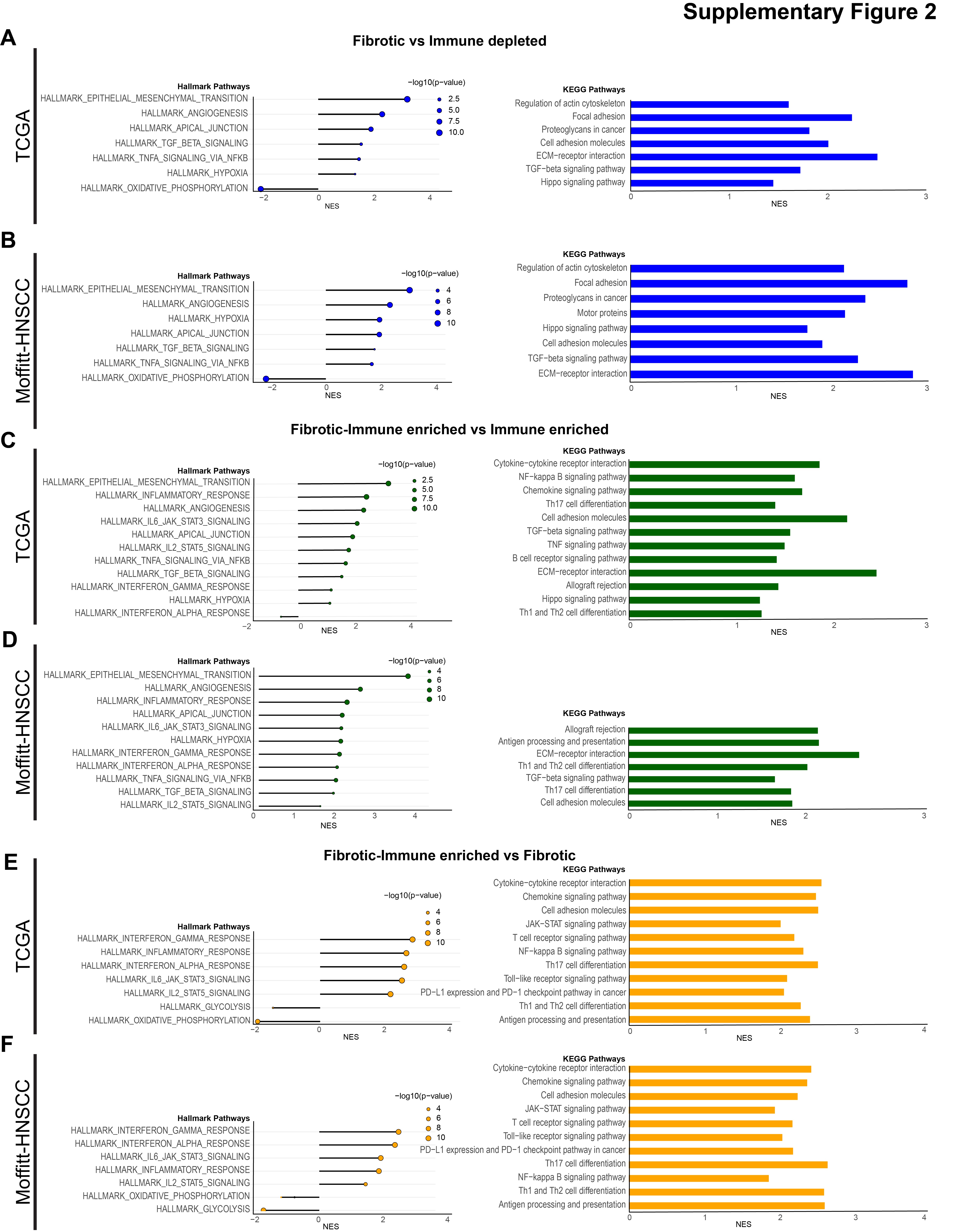

### Supplemental Figure S3

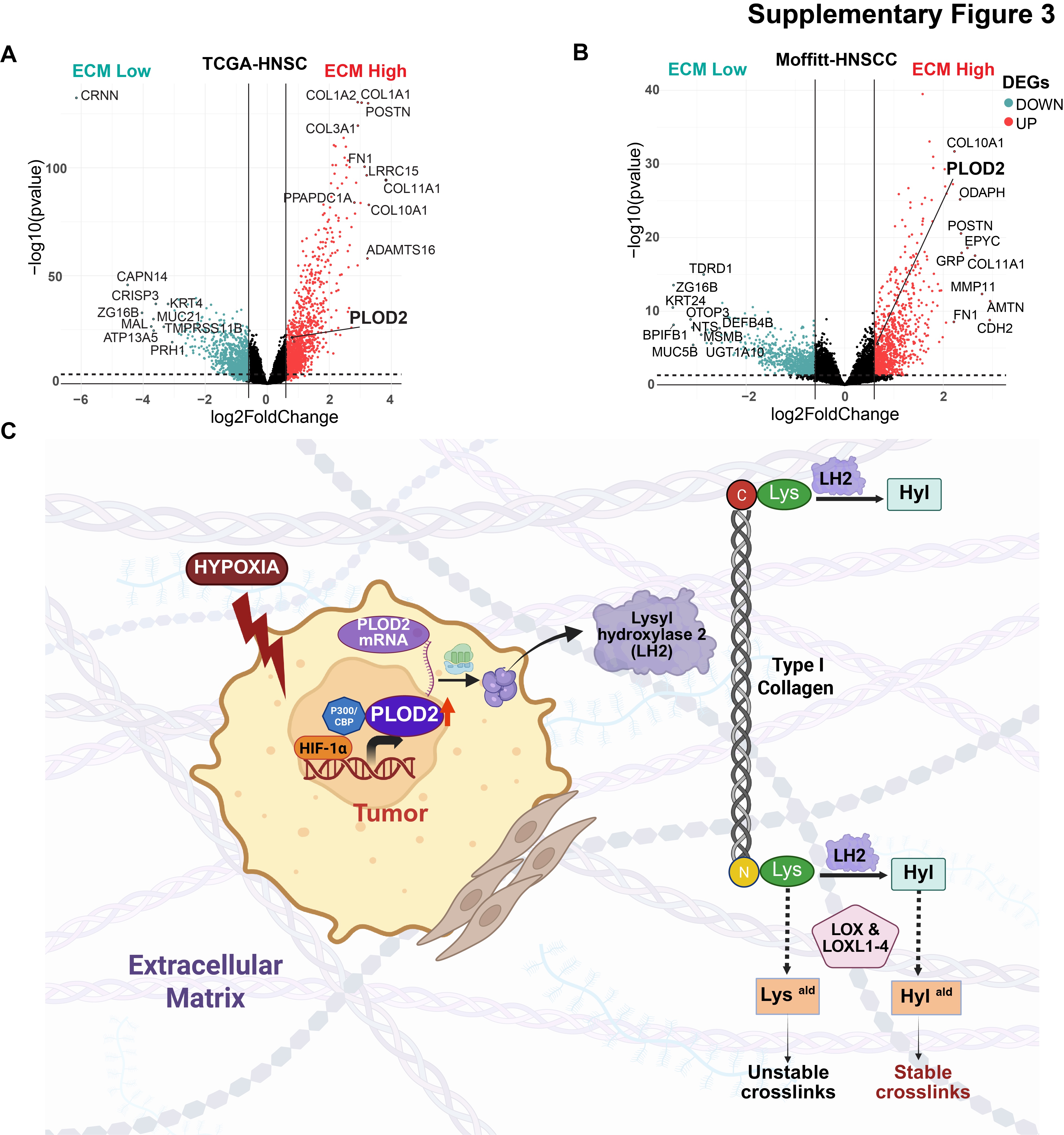

### Supplemental Figure S4

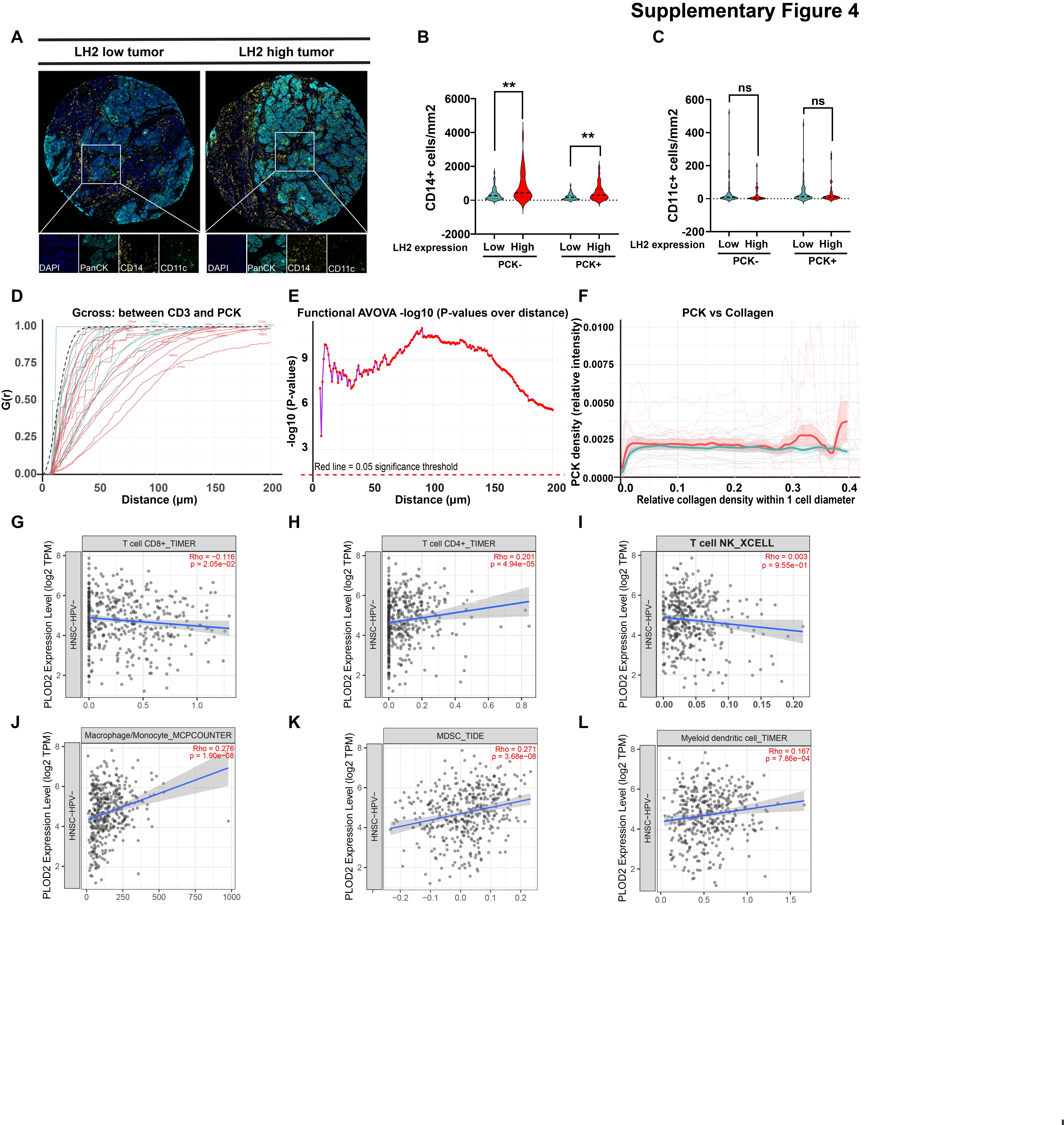

### Supplemental Figure S5

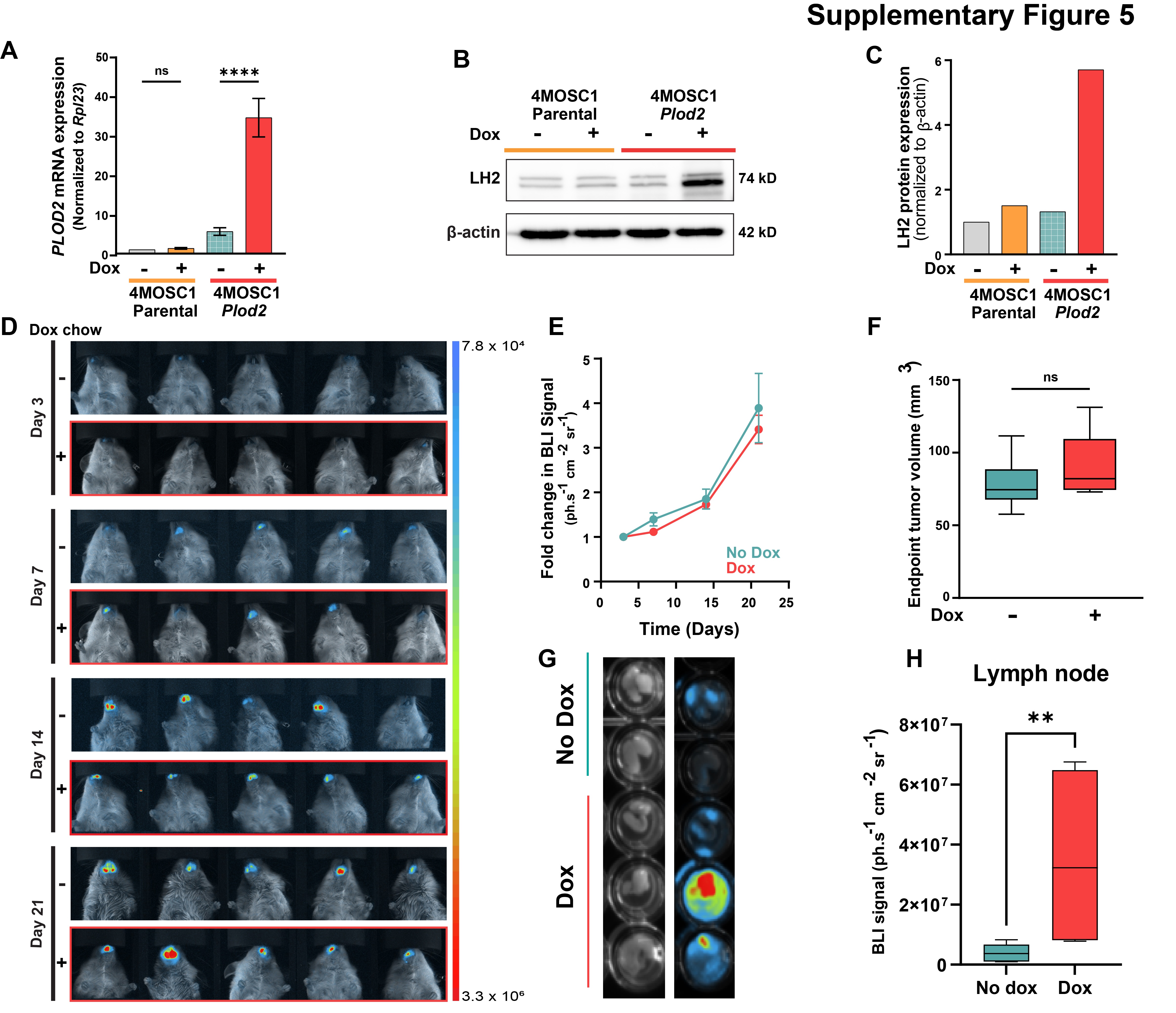

### Supplemental Figure S6

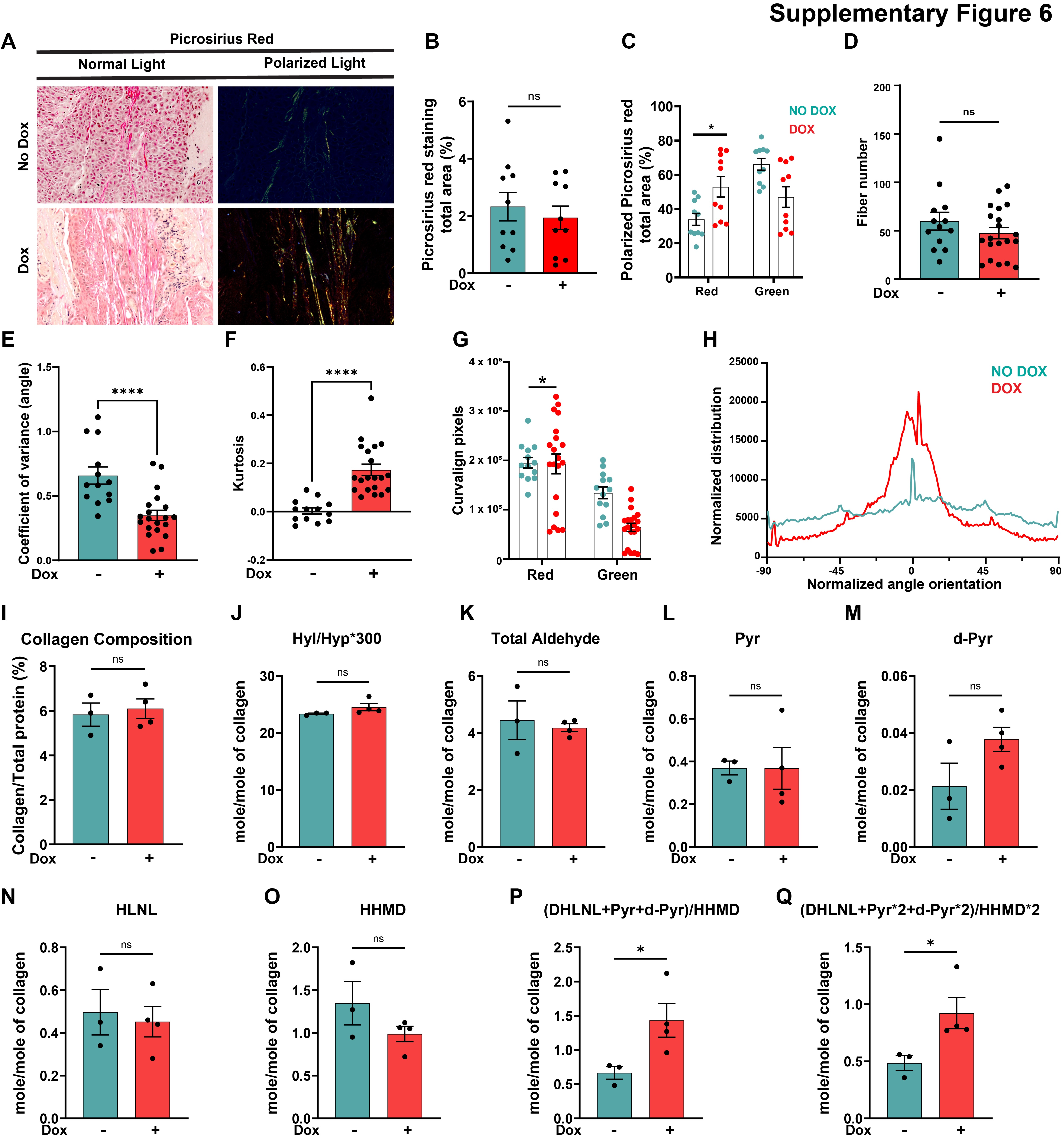

### Supplemental Figure S7

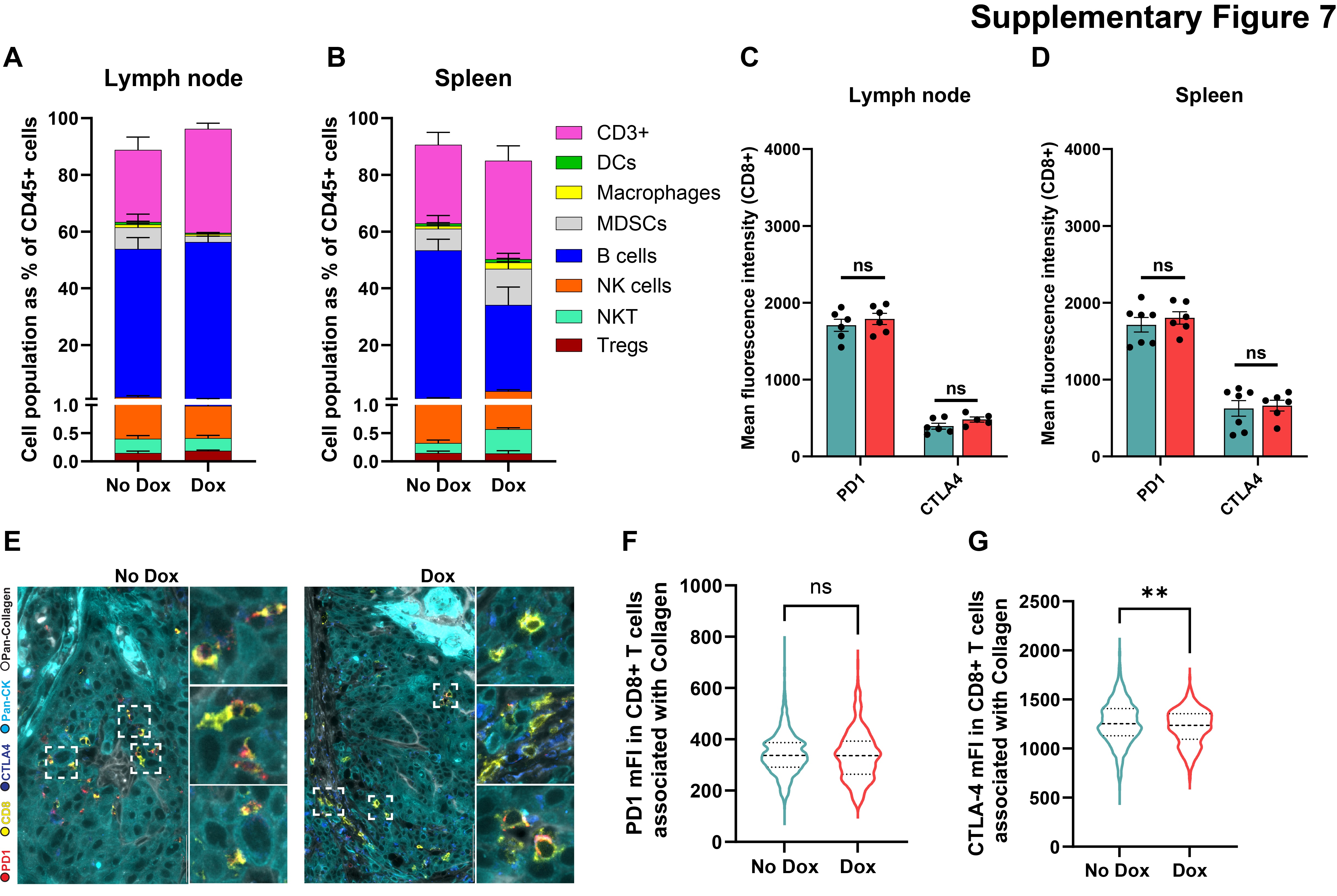

### Supplemental Figure S8

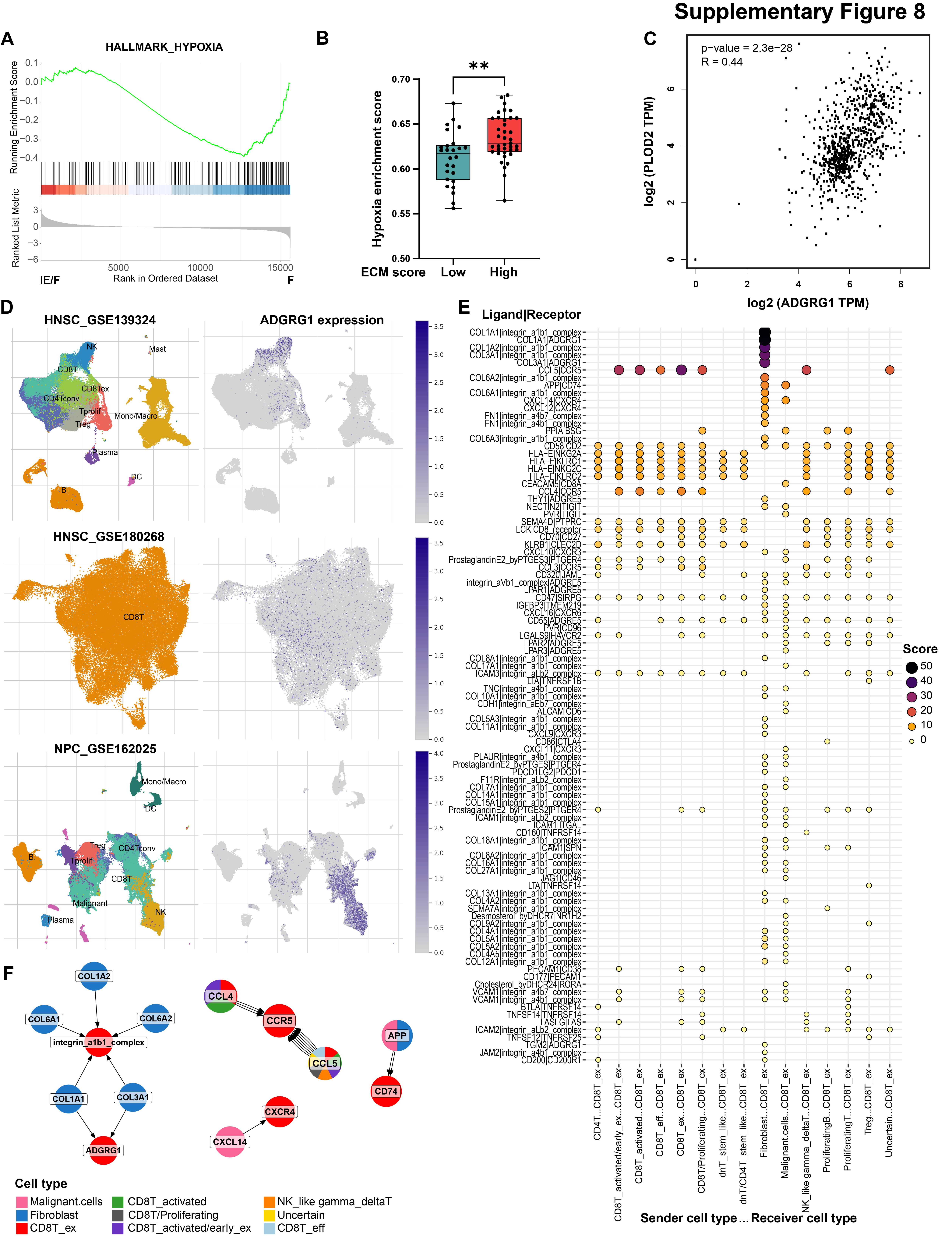

### Supplemental Figure S9

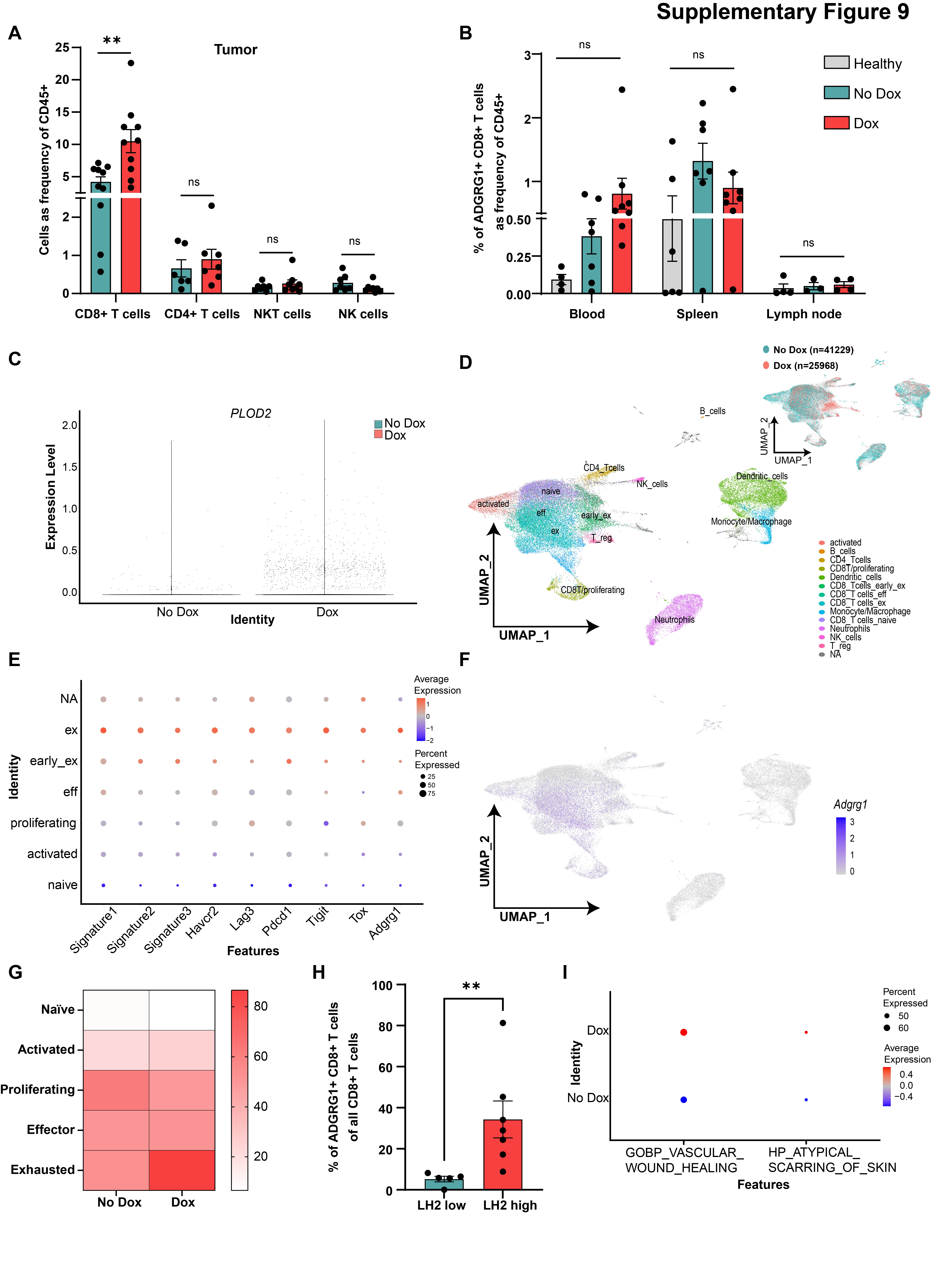

### Supplemental Figure S10

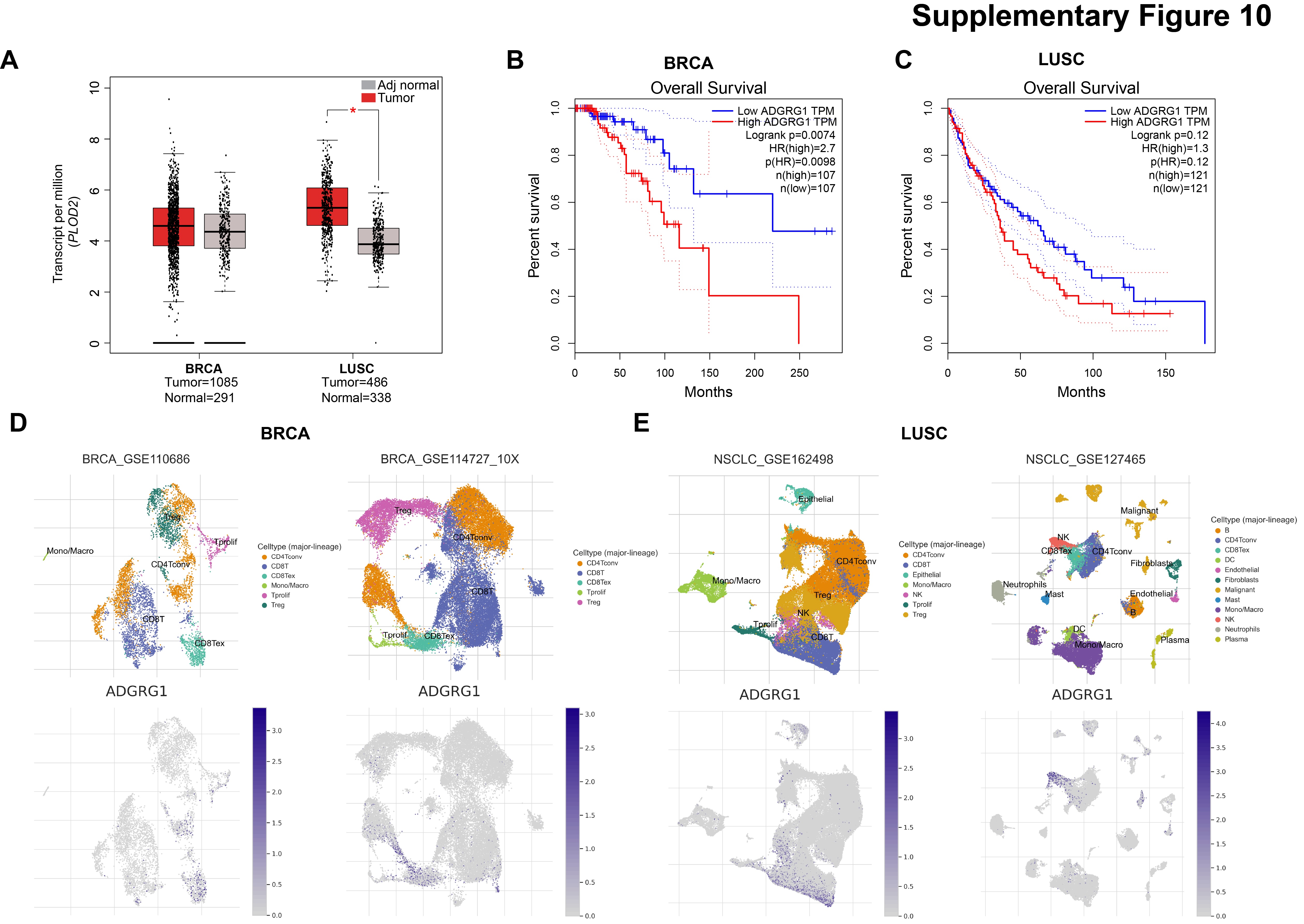

### Supplemental Figure S11

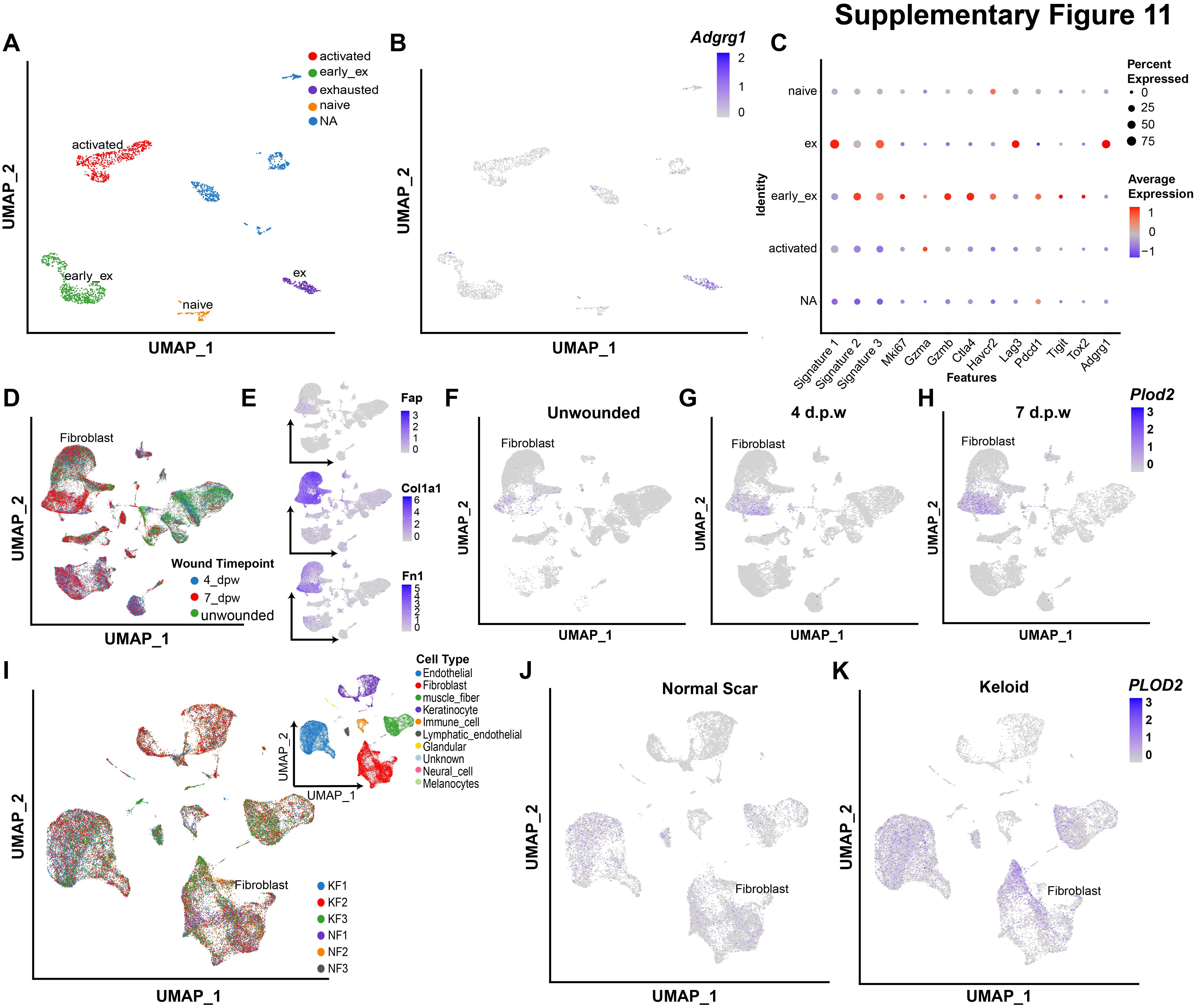

### Supplemental Figure S12

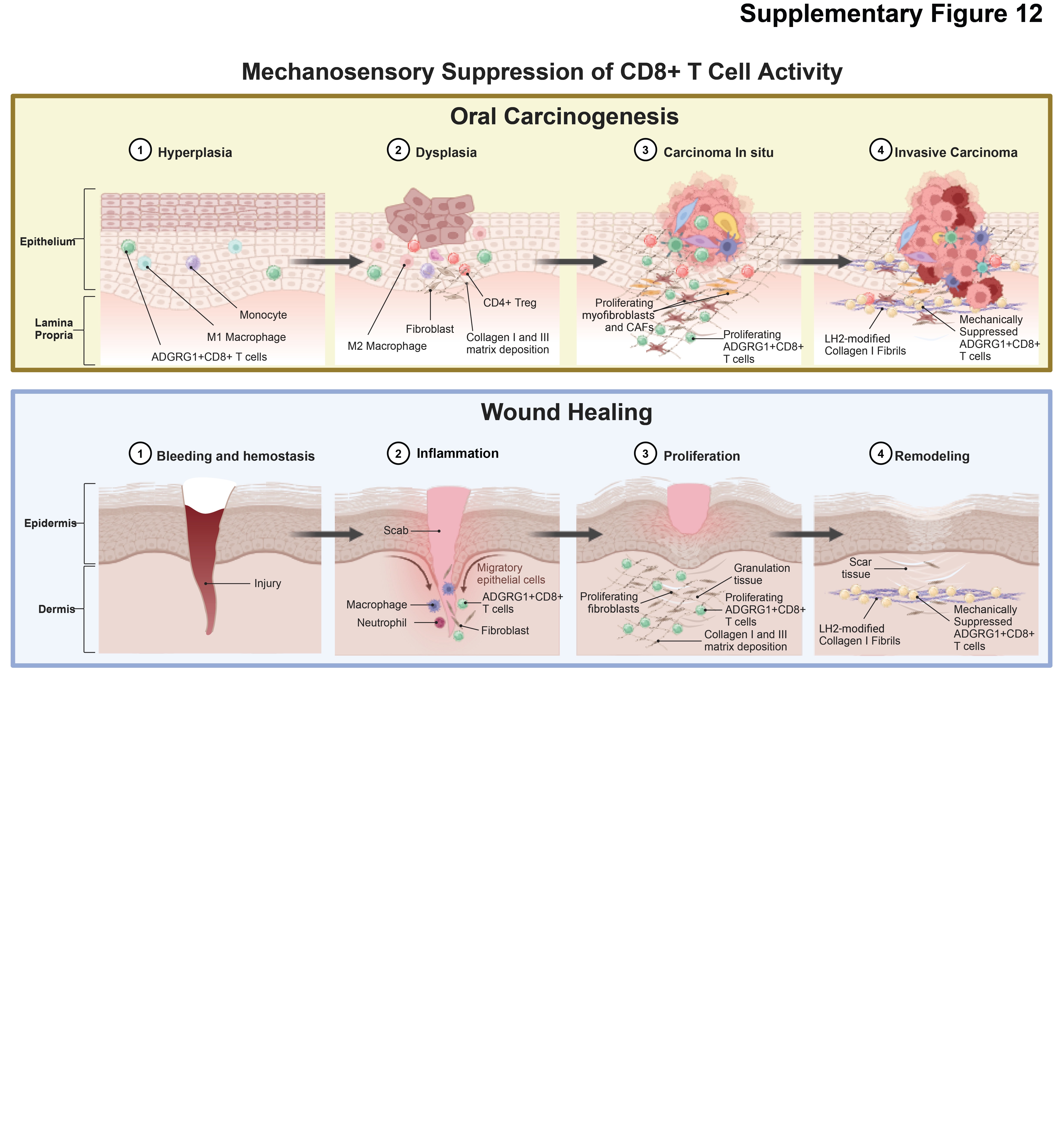
